# Deconvoluting Stress-Responsive Proteostasis Signaling Pathways for Pharmacologic Activation using Targeted RNA-sequencing

**DOI:** 10.1101/499046

**Authors:** Julia M.D. Grandjean, Lars Plate, Richard I. Morimoto, Michael J. Bollong, Evan T. Powers, R. Luke Wiseman

**Affiliations:** Department of Molecular Medicine, The Scripps Research Institute, La Jolla, CA, USA; Department of Chemistry, The Scripps Research Institute, La Jolla, CA, USA; Department of Molecular Biosciences, Rice Institute for Biomedical Research, Northwestern University, Evanston, IL, USA

**Keywords:** Targeted RNAseq, unfolded protein response, heat shock response, oxidative stress response, small molecule activator, screening

## Abstract

Cellular proteostasis is maintained by stress-responsive signaling pathways such as the heat shock response (HSR), the oxidative stress response (OSR), and the unfolded protein response (UPR). Activation of these pathways results in the transcriptional upregulation of select subsets of stress-responsive genes that restore proteostasis and adapt cellular physiology to promote recovery following various types of acute insult. The capacity for these pathways to regulate cellular proteostasis makes them attractive therapeutic targets to correct proteostasis defects associated with diverse diseases. High-throughput screening (HTS) using cell-based reporter assays is highly effective for identifying putative activators of stress-responsive signaling pathways. However, the development of these compounds is hampered by the lack of medium-throughput assays to define compound potency and selectivity for a given pathway. Here, we describe a targeted RNA sequencing (RNAseq) assay that allows cost effective, medium-throughput screening of stress-responsive signaling pathway activation. We demonstrate that this assay allows deconvolution of stress-responsive signaling activated by chemical genetic or pharmacologic agents. Furthermore, we use this assay to define the selectivity of putative OSR and HSR activating compounds previously identified by HTS. Our results demonstrate the potential for integrating this adaptable targeted RNAseq assay into screening programs focused on developing pharmacologic activators of stress-responsive signaling pathways.

## INTRODUCTION

Imbalances in cellular proteostasis can be induced by genetic, environmental, or aging-related insults and are intricately involved in the pathology of multiple, etiologically diverse diseases^1–3^. These include diabetes, cardiovascular disorders, and neurodegenerative diseases such as Alzheimer’s and Parkinson’s disease^1–5^. In order to protect from these types of insults, cells have evolved an integrated network of stress-responsive signaling pathways including the heat shock response (HSR)^6–8^, the oxidative stress response (OSR)^9–11^,the unfolded protein response (UPR)^12–15^, and the integrated stress response (ISR)^16^ (Fig. 1A). These pathways are activated by distinct types of stress and initiate signal transduction pathways that ultimately activate transcription factors such as the HSR-associated Heat Shock Factor 1 (HSF1), the OSR-associated Nuclear Factor Erythroid 2 (NRF2), and the UPR-associated transcription factors X-box Binding Protein 1 (XBP1s), Activating Transcription Factor 6 (ATF6), and Activating Transcription Factor 4 (ATF4) (the latter also being implicated in the ISR)^6–8, 12, 17, 18^. These transcription factors induce select subsets of stress-responsive genes to alleviate specific types of proteostasis stress and promote cellular recovery following an acute insult. The capacity for these signaling pathways to protect cells against different types of proteostasis-related stress makes them highly attractive therapeutic targets to ameliorate pathologic imbalances in proteostasis associated with diverse human diseases^1, 19–24^.

**Figure 1.**
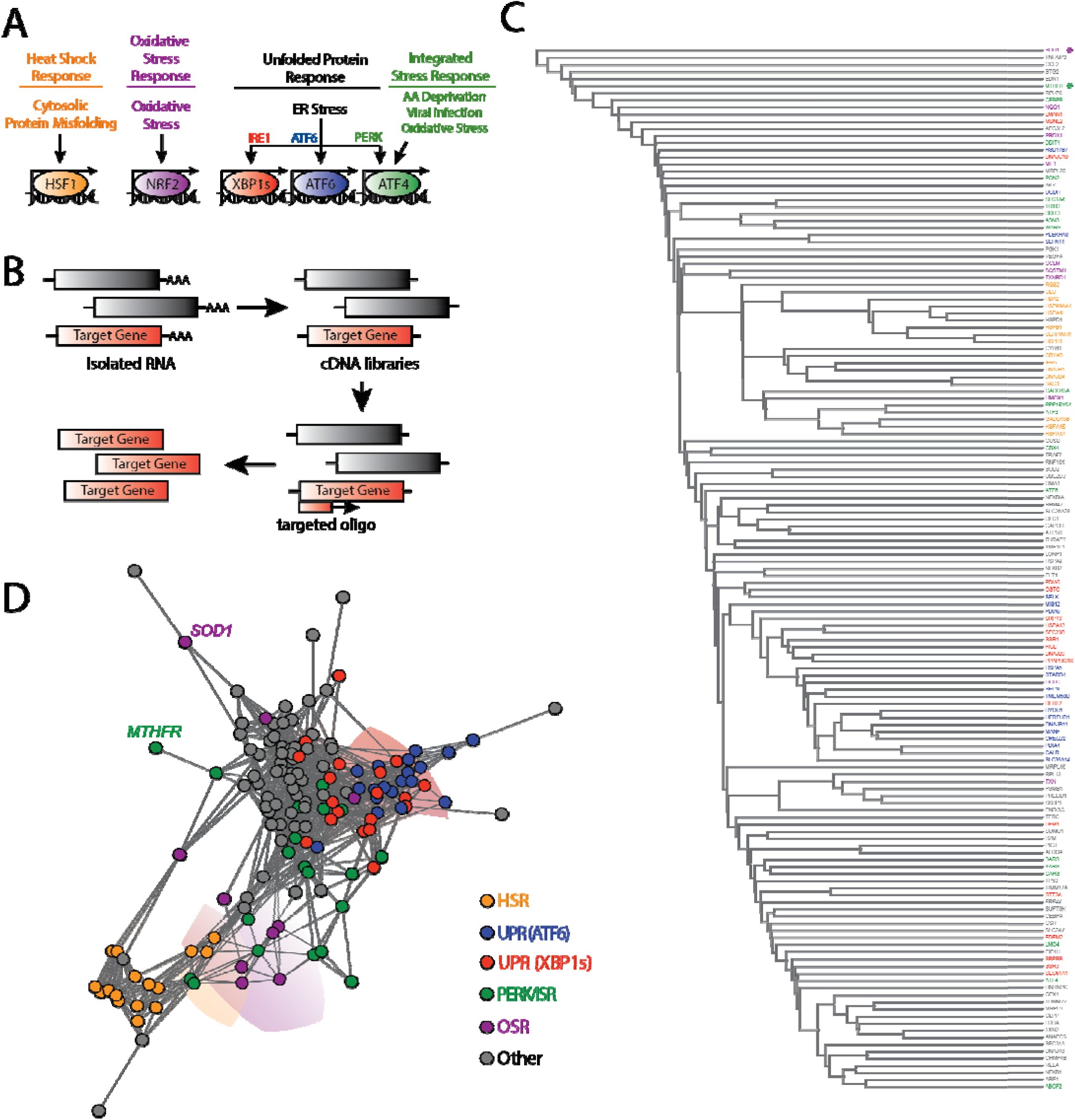
Targeted RNA sequencing deconvolutes stress-responsive transcriptional programs. A. Illustration showing the stress-responsive proteostasis pathways profiled in our targeted RNAseq assay. Stresses that activate each pathway and specific transcription factors activated downstream of these pathways are also shown.
B. Schematic of the general protocol used for our targeted RNAseq assay. Briefly, RNA is isolated from cells following a given treatment. This RNA is then converted into cDNA libraries that are probed using oligos targeted to specific stress-responsive genes (red) for sequencing library generation.
C. Dendrogram of individual target genes from our targeted RNAseq panel (see Table 1) grouped by hierarchical clustering using the Euclidean distance between each genes’ expression level correlation coefficients over all treatment conditions (see Table 2 and **Table S1**). Genes are colored by assignment to specific stress-responsive signaling pathways to report on activation of the HSR (orange), the OSR (purple), the ATF6 UPR signaling pathway (blue), the IRE1/XBP1s UPR signaling pathway (red), the PERK/ISR signaling pathway (green), or other pathways (grey). The asterisks identify *SOD1* (purple) and *MTHFR* (green).
D. Network graph of individual target genes from our targeted RNAseq panel showing the clustering of genes into defined stress-responsive signaling pathways. This graph is derived by representing each gene as a vertex and connecting the vertices for genes whose expression level changes correlate with Pearson *R* > 0.6. Genes that do not correlate at this level with any other genes are connected only to the gene with which they have the highest correlation coefficient. Pathways are colored using the same scheme described above in Fig. 1C. *SOD1* and *MTHFR* are identified by name.

The development of pharmacologic activators of stress-responsive signaling pathways has primarily been pursued using high-throughput screening (HTS) approaches that employ cellular transcriptional reporters of target genes activated downstream of specific stress pathways including the ATF6 signaling arm of the UPR and the HSF1-dependent HSR^20, 25–30^. While this approach has effectively identified many putative activators of these pathways, the further development and characterization of these HTS hits is often hampered by complications including reporter interference, lack of compound selectivity for a given pathway, or reporter constructs not reliably reporting on activation of the entire protective transcriptional program^24, 28, 31, 32^. Without proper tools to assess selectivity across broad stress signaling pathways, it is difficult to determine whether previous HTS have identified effective compounds that selectively activate these pathways.

One strategy to increase the efficiency of identifying specific pathway activators from many screening hits is to incorporate upstream transcriptional profiling to first define the activation spectrum among stress responsive signaling pathways. The benefits of this approach have been demonstrated with the recent establishment of compounds that preferentially activate the ATF6 signaling arm of the UPR, where multiplex gene expression (MGE) profiling was integrated into a screening pipeline centered on cell-based transcriptional reporters^25^. However, despite the evidence highlighting the benefit of incorporating transcriptional profiling into screening platforms, cost effective strategies to profile stress-responsive signaling pathway activation in a medium-throughput format are currently lacking.

Defining the magnitude and repertoire of activation among stress-responsive signaling pathways for a given stimulus is complicated by multiple challenges. Stress-responsive genes can be regulated by multiple signaling pathways, making it difficult to discern pathway activation by tracking the expression of a single gene. For example, the OSR target gene *HMOX1* can be regulated by multiple stress-responsive transcription factors including NRF2 (OSR), HSF1 (HSR), and NF-kB^33^. Furthermore, many stress-responsive signaling pathways have overlapping sets of target genes, challenging the ability to define selective activation of a certain pathway. For example, the majority of genes regulated by the UPR-associated transcription factor ATF6 are also activated, albeit to lower extents, by the alternative UPR-associated transcription factor XBP1s, thus making it difficult to deconvolute specific activation of these pathways by monitoring expression of a single gene^34^. Additionally, different stress-responsive signaling pathways induce target genes to varying extents. For example, HSF1 (HSR) target genes can be induced >10-fold higher than UPR target genes^17, 34^. These properties of stress-responsive signaling challenge the ability to monitor activation of specific pathways using strategies such as geneset enrichment analysis (GSEA), which is biased towards pathways that elicit larger transcriptional responses and does not easily deconvolute overlapping stress-responsive transcriptional programs. Furthermore, GSEA requires whole transcriptome RNA sequencing (RNAseq) profiling to define pathway activation, limiting its application as a medium-throughput screening approach. One potential strategy to address the above challenges is to monitor activation of specified sets of stress-responsive genes regulated downstream of different stress-responsive signaling pathways, wherein pathway activation is defined by the grouped behavior for all relevant target genes. This strategy requires measuring multiple genes activated downstream of different stress-responsive signaling pathways in a cost-effective assay.

Recent advances in RNA sequencing have demonstrated the potential for this approach to be integrated into drug discovery pipelines. For example, the DRUG-seq platform established a cost-effective strategy to profile compounds in a high-throughput format, providing a powerful approach to define compound mechanism of activation and selectivity^35^. However, this approach requires specialized equipment that would make it difficult to implement in most research laboratories. In contrast, the targeted RNAseq platform, described in the last 5 years (previously described as Capture-seq^36^), provides a unique opportunity to screen expression of 100–1000 genes in a cost-effective, medium-throughput format. Since this approach uses target-specific generation of sequencing libraries, targeted RNAseq demonstrates improved sensitivity for low-copy transcripts, potentially providing a larger dynamic range to track changes in both low and highly-expressed genes. Furthermore, targeted RNAseq avoids background issues caused by non-specific probe binding or probe cross-hybridization found in such technologies as microarrays^37^. As such, this approach has been used in diverse contexts including measuring expression of alleles in plant populations^38^, detection of gene fusions in solid tumors^39^, and monitoring activation of cell death pathways^40^.

Here, we describe a targeted RNAseq assay designed to define activation of stress-responsive proteostasis pathways in a medium-throughput format. We show that this approach allows accurate deconvolution of stress-responsive pathway activation induced by diverse chemical genetic and pharmacologic agents. Furthermore, we demonstrate the potential for this approach to define the selectivity of pharmacologic activators of stress-responsive signaling pathways by profiling the selectivity of compounds identified by high-throughput reporter screening to activate the OSR-associated transcription factor NRF2 or the HSR-associated transcription factor HSF1^20, 26^. Ultimately, our results show that targeted RNAseq profiling is a highly adaptable strategy that can be efficiently incorporated into HTS pipelines and downstream compound development to improve the establishment of pharmacologic activators of specific stress-responsive signaling pathways.

## RESULTS & DISCUSSION

### A Targeted RNAseq Assay to Monitor Activation of Stress-Responsive Proteostasis Pathways

To establish a targeted RNAseq assay for monitoring activation of stress-responsive proteostasis pathways, we first defined genesets predicted to accurately report on the activation of the predominant proteostasis pathways: the HSR, OSR, the three signaling arms of the UPR, and the ISR (Fig. 1A). We used published transcriptional profiles from human cells subjected to pharmacologic, chemical genetic, and genetic manipulation of specific stress-responsive signaling pathways to identify sets of proteostasis genes induced by each pathway. From this data, we selected 10–20 reporter genes activated downstream of the HSR^17^, OSR^41^, and the three arms of the UPR regulated by the ER stress sensing proteins IRE1, ATF6, and PERK^34, 42–44^ (Fig. 1A and Table 1). Importantly, the geneset that reports on activation of the PERK signaling arm of the UPR can also be used to monitor activation of the ISR, as both are activated through a process involving phosphorylation of the α subunit of eukaryotic initiation factor 2 (eIF2α) and the activity of the ATF4 transcription factor^18, 44^. In addition, we include stress-responsive genes activated downstream of other stress-responsive signaling pathways including the Hypoxic Stress Response^45^, NFκB signaling^41, 46^, and the poorly defined mitochondrial unfolded protein response (UPR^mt^)^47–49^, providing additional readouts of stress-pathway activation. Our final gene panel consists of 150 target genes shown in Table 1.

**Table 1.**
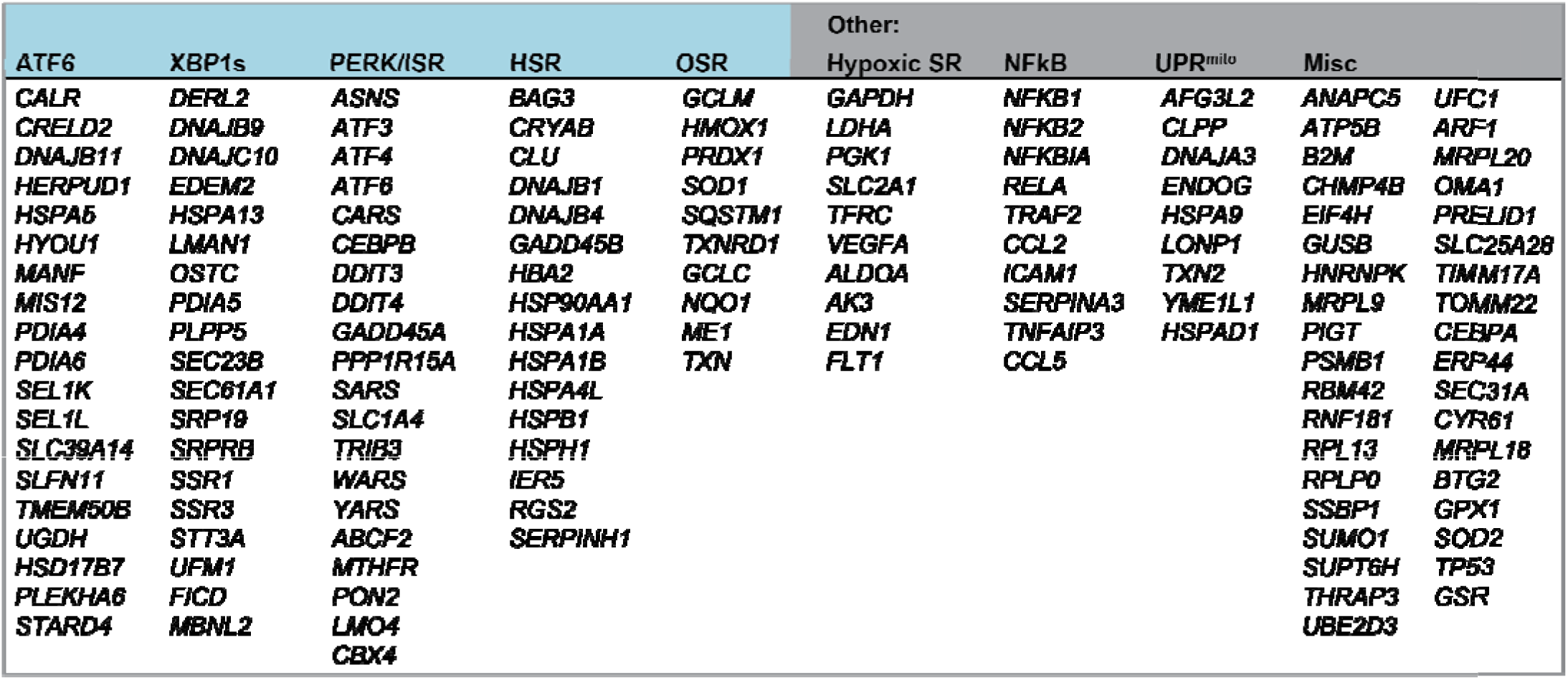
Targeted RNAseq gene panel to report on activation of stress-responsive proteostasis pathways

We used the established targeted RNAseq profiling approach with this custom gene panel to define the activation of stress-responsive signaling pathways in HEK293T cells grown in 96-well plates subjected to multiple conditions predicted to activate the different stress-responsive proteostasis pathways shown in Fig. 1A (see Table 2 and **Table S1**). Briefly, we isolated RNA from these cells and generated cDNA libraries using a standard reverse transcriptase reaction. We then amplified our genes of interest for sequencing using targeted primer sets directed to the 150 genes in the panel (Fig. 1B). Amplicons were then isolated and pooled for sequencing using the MiSeq desktop sequencer at a target depth of 50 million paired-end reads for all pooled samples. Overall alignment of reads reflected the specific nature of this approach with over 93% of reads aligning to target regions, which is significantly greater than that observed in conventional whole transcriptome RNAseq experiments (**Fig. S1A**). Our desktop MiSeq sequencing run yielded a median of ~580,000 reads per sample (19 conditions in triplicate, 57 samples total), which is approximately 1% of the number of reads aligned per sample with whole transcriptome RNAseq on the same platform (approximated to 44–50×10^6^ reads per sample). Additionally, per gene target, our targeted RNaseq assay yielded an median of 721.7 aligned reads (median total >41,000 reads) across all treatment conditions included in these analyses (**Fig. S1B-C**). All of the data from this targeted RNAseq assay is included in **Table S2**.

**Table 2.** Treatment conditions for Stress Signaling Targeted RNAseq

| Cell Type | Treatment | Targeted Pathway |
| --- | --- | --- |
| 293TcHSF1 | arsenite (As(III)) | HSR/OSR/ISR |
| 293TcHSF1 | dox-HSF1 | HSR |
| 293<sup>DAX</sup> | dox-XBP1s | XBP1s |
| 293<sup>DAX</sup> | DHFR.ATF6 | ATF6 |
| 293<sup>DAX</sup> | dox-XBP1s+DHFR.ATF6 | XBP1s+ATF6 |
| 293<sup>DAX</sup> | thapsigargin (Tg) | UPR |
| 293T | thapsigargin (Tg) | UPR |
| 293T | tunicamycin (Tm) | UPR |
| 293T | oligomycin | ISR/OSR |
| 293T | paraquat | ISR/OSR |
| 293T | CBR-470-1 | OSR |
| 293T | bardoxolone | OSR |
| 293T | A3 | HSR |
| 293T | C1 | HSR |
| 293T | D1 | HSR |
| 293T | F1 | HSR |

Across replicate samples, we observed high reproducibility with all treatments demonstrating an R^2^-value above 0.7 and most having an R^2^>0.85 (**Fig. S1D-E**). From aligned count data, we performed unbiased clustering across all treatment conditions to determine our ability to accurately define different stress-signaling pathways (Fig. 1C,D). This analysis shows that genes regulated by the HSR, OSR, and the three arms of the UPR (IRE1/XBP1s, ATF6 and PERK/ISR) generally cluster together, reflecting their similar regulation across the various stress conditions. However, despite this clustering, we observe significant overlap between genesets, reflecting their integrated activation in response to diverse types of stimuli. For example, the IRE1/XBP1s (red in Fig. 1D) and ATF6 (blue in Fig. 1D) genesets show significant overlap, reflecting the coordinated activation of these two pathways in response to ER stress. Furthermore, certain genes such *as SOD1*, activated downstream of the OSR-associated transcription factor NRF2^50^, separate from the OSR cluster (purple in Fig. 1C-D), reflecting the ability for this gene to be regulated by multiple stress-responsive signaling pathways apart from the OSR^51^. The PERK/ISR target *MTHFR* is also regulated by other UPR signaling pathways, as well as OSR^52, 53^, and is similarly found to separate from the larger cluster of PERK/ISR targets (green in Fig. 1C-D). The overlap of genesets and promiscuity for specific genes to report on multiple pathways highlights the importance of tracking sets of stress-responsive genes for defining overall pathway activation. However, the general clustering of our stress-pathway genesets indicates that this targeted RNAseq assay is capable of tracking changes in stress responsive genes to accurately define activation of specific stress-responsive signaling pathways.

### Defining stress-independent HSR and UPR signaling pathways through targeted RNAseq profiling

We initially validated the ability for our targeted RNAseq assay to report on activation of specific stress-signaling proteostasis pathways using chemical genetic approaches that allow activation of specific pathways independent of stress. First, we defined activation of the HSR-regulated proteostasis genes in HEK293^trex^ cells following doxycycline (dox)-dependent activation of a constitutively active cHSF1^17^ – the primary transcription factor regulated by the HSR^6–8^. In order to define the induction of specific target proteostasis genes in our targeted RNAseq data, we first median-normalized aligned counts per target gene across all treatment conditions. We took average normalized count values across sample replicates and performed a Log_2_ transformation, yielding the “Log_2_ normalized counts” used for relative expression analysis. To compare chemical genetic and pharmacologic activating conditions versus vehicle control samples, we conducted a correlation analysis of Log_2_ normalized counts to yield a line of best fit (Fig. 2A), reflecting baseline expression levels for the majority of genes not affected by a given treatment. We then calculated the deviation of each target gene for the experimental condition from the line of best fit, or “residual value”, which was used to quantify up/downregulation of that gene. (Fig. 2A). Finally, we define pathway activation by plotting the residual values of each gene from this analysis, grouped according to assigned stress-responsive pathway, and monitoring the overall behavior of the geneset (Fig. 2B). This allows us to normalize variability in gene induction across different treatments and minimize challenges associated with lowly expressed genes that show high levels of induction. From this analysis, we demonstrate that dox-dependent cHSF1 activation robustly and selectively activates the entire target HSR-regulated proteostasis program, thus confirming the ability for our targeted RNAseq assay to define activation of this pathway (Fig. 2B and **Table S3**). Interestingly, the activation of this pathway is identical to that observed when we perform the same analysis using published RNAseq transcriptional profiles for dox-dependent cHSF1 activation^17^, demonstrating that our RNAseq assay accurately quantifies the induction of HSR-regulated proteostasis target genes (**Fig. S2A-C**).

**Figure 2.**
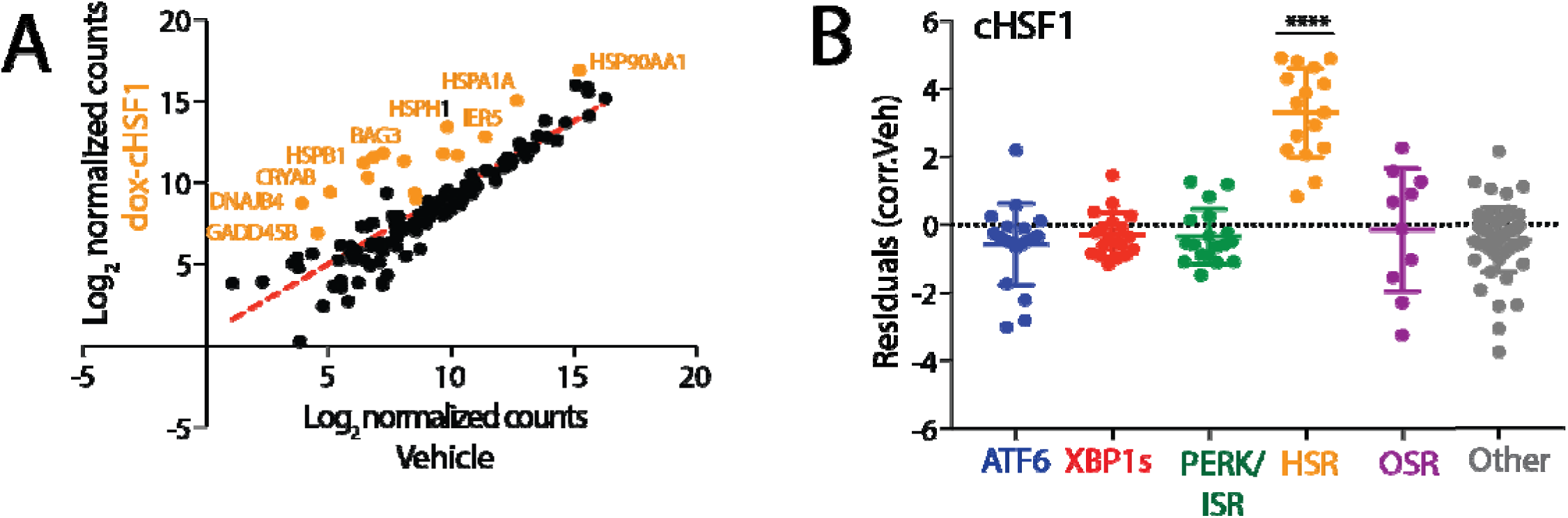
Targeted RNAseq accurately defines HSR activation induced by stress-independent activation of cHSF1. A. Log_2_ normalized aligned transcript counts for HEK293^trex^ cells expressing doxycycline (dox)-inducible cHSF1 treated with 2.25 μM dox (y-axis) or vehicle (x-axis) for 16 h. Aligned transcript counts represent averages from three independent replicates quantified from our targeted RNAseq assay. All identified genes are HSR target genes.
B. Plot showing residuals calculated by comparing the expression of our panel of stress-responsive genes between HEK293^trex^ cells expressing dox-inducible cHSF1 following 16 h treatment with dox (2.25 μM) or vehicle. Calculation of residuals was performed as described in Fig. 2A. Statistics were calculated using one-way ANOVA. Significance shown reflects comparison to “Other” target transcript set. **p<0.01, ***p<0.001, ****p<0.0001. See **Table S3** for full ANOVA table.

A significant challenge in monitoring activation of stress-responsive signaling pathways is the overlap between closely related pathways. For example, the IRE1/XBP1s UPR pathway induces expression of multiple genes also regulated by the ATF6 UPR signaling pathway, albeit to lower extents^34^. Furthermore, other XBP1s target genes are often induced to lower levels than that observed for ATF6-selective target genes^34^. To define the potential for targeted RNAseq to discern selective activation of these two UPR signaling pathways, we performed this assay in HEK293^DAX^ cells subjected to stress-independent XBP1s and/or ATF6 activation. HEK293^dax^ cells express dox-inducible XBP1s and trimethoprim (TMP)-regulated DHFR.ATF6, allowing activation of these two transcription factors in the same cell independent of ER stress through administration of dox and/or TMP^34^. As a control, we also monitored gene expression in response to global ER stress induced by treating HEK293^DAX^ cells with the SERCA inhibitor, thapsigargin (Tg). As expected, Tg treatment showed robust activation of all three UPR signaling pathways (IRE1/XBP1s, ATF6, and PERK/ISR), confirming global UPR activation (**Fig. S3A-B**). In contrast, TMP-dependent DHFR-ATF6 activation showed significant increases in the ATF6 target geneset, consistent with selective ATF6 activation (Fig. 3A). However, dox-inducible XBP1s increased expression of both the IRE1/XBP1s and ATF6 target genesets, although ATF6 target genes were induced less than that observed following ATF6 activation (Fig. 3B), which is consistent with previous work^34^. Combined treatment with a combination of dox (activated XBP1s) and TMP (activated DHFR.ATF6) elicited a strong upregulation of both genesets (Fig. 3C).

**Figure 3.**
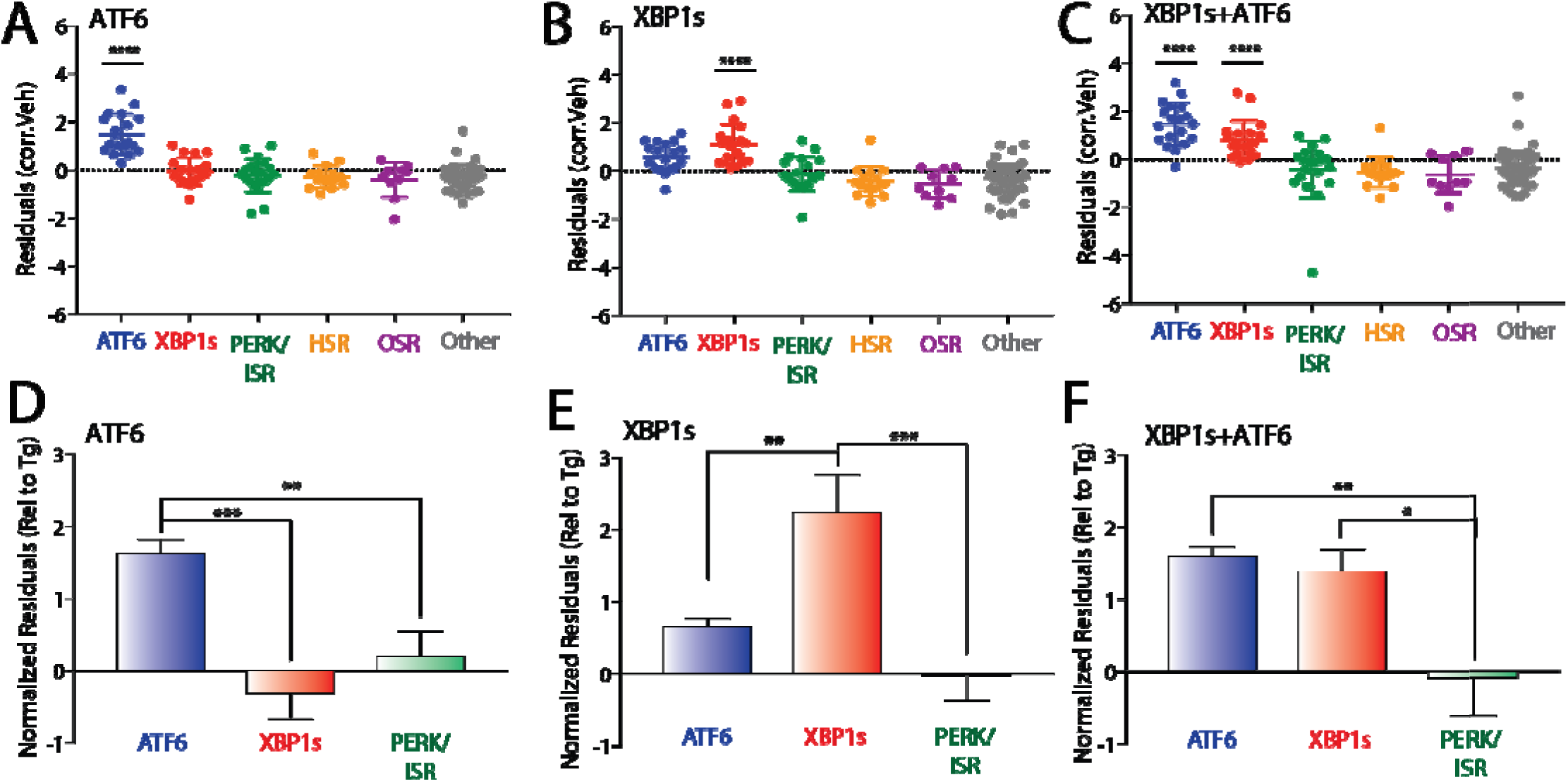
Targeted RNAseq defines stress-independent activation of UPR-associated signaling pathways. A. Plot showing residuals calculated by comparing the expression of our panel of stress-responsive genes between HEK293^dax^ cells following treatment with trimethoprim (10 μM 4 h; activates DHFR.ATF6) or vehicle. Calculation of residuals was performed as described in Fig. 2A. Statistics were calculated using one-way ANOVA. Significance shown reflects comparison to “Other” target transcript set. ****p<0.0001. See **Table S3** for full ANOVA table.
B. Plot showing residuals calculated by comparing the expression of our panel of stress-responsive genes between HEK293^dax^ cells following treatment with dox (1 μg/mL 4 h; activates dox-inducible XBP1s) or vehicle. Calculation of residuals was performed as described in Fig. 2A. Statistics were calculated using one-way ANOVA. Significance shown reflects comparison to “Other” target transcript set. ****p<0.0001. See **Table S3** for full ANOVA table.
C. Plot showing residuals calculated by comparing the expression of our panel of stress-responsive genes between HEK293^dax^ cells following treatment with both trimethoprim (10 μM, 4 h; activates DHFR.ATF6) and dox (1 μg/mL 4 h; activates dox-inducible XBP1s) or vehicle. Calculation of residuals was performed as described in Fig. 2A. Statistics were calculated using one-way ANOVA. Significance shown reflects comparison to “Other” target transcript set. ****p<0.0001. See **Table S3** for full ANOVA table.
D. Graph showing normalized residuals for genesets regulated by the ATF6 (blue), XBP1s (red) or PERK (green) UPR signaling pathways in HEK293^dax^ cells following treatment with TMP (10 μM, 4 h; activates DHFR.ATF6). The residuals for each gene were normalized to those observed for thapsigargin (Tg)-induced ER stress in HEK293^dax^ cells (**Fig. S3A,B**). Normalized data was subjected to ROUT outlier testing. Statistics from one-way ANOVA **p<0.01, ***p<0.001.
E. Graph showing normalized residuals for genesets regulated by the ATF6 (blue), XBP1s (red) or PERK (green) UPR signaling pathways in HEK293^dax^ cells following treatment with dox (1μg/mL, 4 h; activates dox-inducible XBP1s). The residuals for each gene were normalized to those observed for thapsigargin (Tg)-induced ER stress in HEK293^dax^ cells (**Fig. S3A,B**). Normalized data was subjected to ROUT outlier testing. Statistics from one-way ANOVA **p<0.01, ***p<0.001.
F. Graph showing normalized residuals for genesets regulated by the ATF6 (blue), XBP1s (red) or PERK (green) UPR signaling pathways in HEK293^dax^ cells following treatment with both TMP (10 μM, 4 h; activates DHFR.ATF6) and dox (1μg/mL, 4 h; activates dox-inducible XBP1s). The residuals for each gene were normalized to those observed for thapsigargin (Tg)-induced ER stress in HEK293^dax^ cells (**Fig. S3A,B**). Normalized data was subjected to ROUT outlier testing. Statistics from one-way ANOVA. *p<0.05, **p<0.01.

Previous reports indicate that the overlap between XBP1s and ATF6 target gene expression observed following stress-independent activation could be deconvoluted by normalizing the expression of genes to that observed with Tg treatment, providing a way to sensitively define the extent of pathway activation^25^. Performing this normalization shows that TMP-dependent DHFR.ATF6 activation selectively induces expression of ATF6 target genes to levels similar to those observed for Tg-dependent ER stress (Fig. 3D). Importantly, dox-dependent XBP1s activation selectively induces expression of IRE1/XBP1s target genes to levels similar to that observed with Tg by this analysis, while only moderately affecting ATF6 target gene expression (Fig. 3E). This profile is distinct from that observed in cells where XBP1s and ATF6 are co-activated, which shows significantly higher induction of both genesets (Fig. 3F). Importantly, when residual values from our targeted RNAseq analysis are compared to transcriptional changes from whole-transcriptome RNAseq collected from HEK293^dax^ cells subjected to XBP1s and/or ATF6 activation, there is a clear correlation between the two data sets, especially in upregulated targets (**Fig. S3C-E**). These results demonstrate that our targeted RNAseq assay can sensitively deconvolute the complex integration of stress-responsive signaling pathways involved in UPR signaling.

### Targeted RNAseq profiling defines stress-pathway activation induced by cellular toxins

We next used our targeted RNAseq assay to profile activation of stress-responsive signaling pathways induced by chemical toxins including tunicamycin (Tm; an ER stressor that inhibits N-linked glycosylation), the environmental toxin arsenite (As(III)), the mitochondrial ATP synthase inhibitor oligomycin, and the ROS generating compound paraquat (PQ). As predicted, our assay demonstrates that these compounds induce unique activation profiles of different stress-responsive signaling pathways. Consistent with the selective induction of ER stress, Tm treatment activates the three arms of the UPR without globally impacting other stress-responsive signaling pathways (Fig. 4A). In contrast, As(III) induces robust activation of the cytosolic HSR, OSR, and ISR signaling pathways (Fig. 4B), highlighting the promiscuous nature of this toxin for cytosolic proteostasis pathway activation^54^. Oligomycin treatment only significantly activated the ISR geneset, reflecting emerging evidence showing that mitochondrial stress promotes signaling through this pathway (Fig. 4C) ^8 48, 49, 55^. PQ treatment also showed modest increases in ISR genes, although the entire pathway was not significantly activated (**Fig. S4**). However, while our genesets report on activation of whole pathways, numerous individual stress-responsive genes from multiple pathways were induced upon these different treatments. For example, the OSR target gene *HMOX1* is induced in cells treated with mitochondrial toxins oligomycin and paraquat, although we do not observe induction of other OSR target genes. Since *HMOX1* can be regulated by multiple stress-responsive signaling pathways^33^, these results suggest that administration of these toxins induce pleiotropic effects on multiple stress-responsive signaling pathways outside of the four primary proteostasis pathways profiled in our targeted RNAseq platform. Regardless, it is clear that our targeted RNAseq assay does accurately reflect predicted toxin-induced activation of proteostasis pathways, further validating the benefit of this approach for profiling pharmacologic activators of stress-responsive proteostasis pathways.

**Figure 4.**
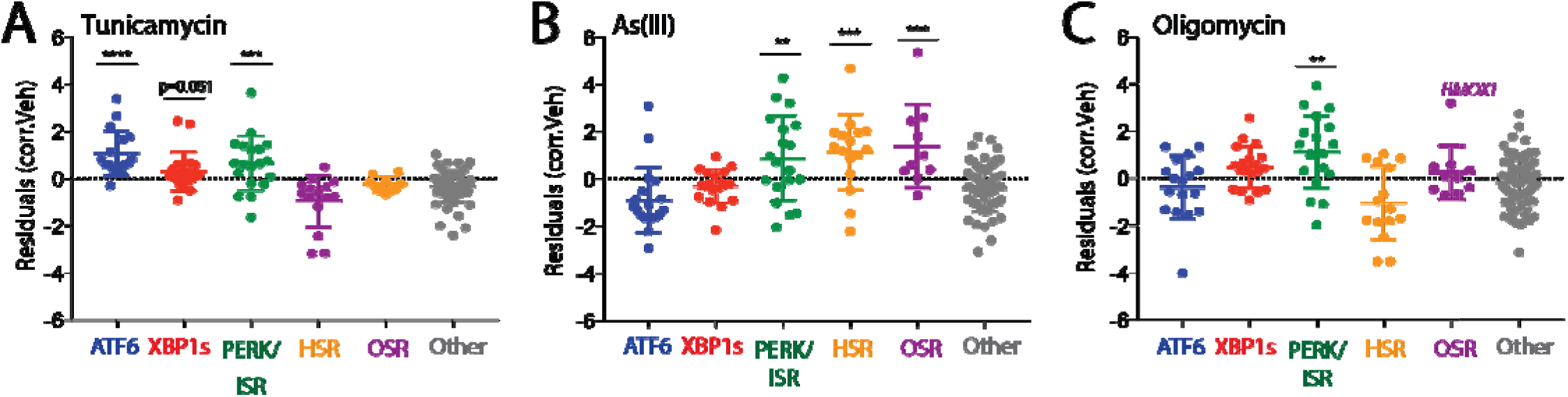
Targeted RNASeq profiling defines activation of stress-responsive signaling pathways induced by diverse environmental toxins. A. Plot showing residuals calculated by comparing the expression of our stress-responsive gene panel between HEK293T cells following treatment with tunicamycin (Tm; 10 μM, 4 h) or vehicle. Calculation of residuals was performed as described in Fig. 2A. Genes are grouped by target stress-responsive signaling pathway. Statistics were calculated using one-way ANOVA, Significance shown reflects comparison to “Other” target transcript set. ***p<0.001, ****p<0.0001. See **Table S3** for full ANOVA table.
B. Plot showing residuals calculated by comparing the expression of our stress-responsive gene panel between HEK293T cells following treatment with arsenite (As(III); 25 μM, 16 h) or vehicle. Calculation of residuals was performed as described in Fig. 2A. Genes are grouped by target stress-responsive signaling pathway. Statistics were calculated using one-way ANOVA, Significance shown reflects comparison to “Other” target transcript set. **p<0.01, ***p<0.001. See **Table S3** for full ANOVA table.
C. Plot showing residuals calculated by comparing the expression of our stress-responsive gene panel between HEK293T cells following treatment with oligomycin A (Oligo; 100 nM, 24 h) or vehicle. Calculation of residuals was performed as described in Fig. 2A. Genes are grouped by target stress-responsive signaling pathway. Statistics were calculated using one-way ANOVA, Significance shown reflects comparison to “Other” target transcript set. **p<0.01. See **Table S3** for full ANOVA table.

### Defining selectivity of pharmacologic NRF2 activating compounds through Targeted RNAseq transcriptional profiling

We next employed our targeted RNAseq assay to define the selectivity of two putative NRF2 activating compounds: bardoxolone and the recently described CBR-470-1 (**Fig. S5A**)^26^. Bardoxolone is an anti-inflammatory compound currently in clinical trials for chronic kidney disease. This compound is reported to induce protective benefits through activation of the OSR-associated transcription factor NRF2^22, 56^. However, it also covalently modifies hundreds of proteins^57^ and displays additional cellular activities including inhibition of the mitochondrial protease LON^58^, suggesting that, apart from NRF2, bardoxolone could also activate other stress-responsive signaling pathways. Interestingly, we show in HEK293T cells that bardoxolone significantly induces expression of the OSR-target gene *HMOX1*, but not other OSR target genes, suggesting that this compound does not robustly activate the NRF2 transcriptional response in these cells (Fig. 5A). However, this compound does induce both the HSR and the ISR genesets, indicating promiscuous activity for this pharmacologic agent. Furthermore, we see strong upregulation of the ATF6 target gene *HSPA5 (also known as BiP)*, without complete activation of the ATF6 pathway. These results indicate that bardoxolone induces pleiotropic effects on stress-responsive genes outside of NRF2 activation. In contrast, CBR-470-1 showed selective activation of the OSR geneset with no significant induction of other stress pathways, suggesting improved selectivity of CBR-470-1 for OSR activation (Fig. 5B). Consistent with this, qPCR analysis of *BAG3* (an HSR target) and *HSPA5* shows that bardoxolone promiscuously induces these non-NRF2 target genes, while CBR-470-1 does not (Fig. 5C,D). However, both compounds induce activation of the OSR target gene *HMOX1* (Fig. 5E). This result is identical to that observed by our targeted RNAseq analysis (Fig. S5B-D). These results show that CBR-470-1 shows increased selectivity for OSR activation relative to bardoxolone and demonstrates the utility for our targeted RNAseq assay to profile selectivity of putative OSR activating compounds in clinical development.

**Figure 5.**
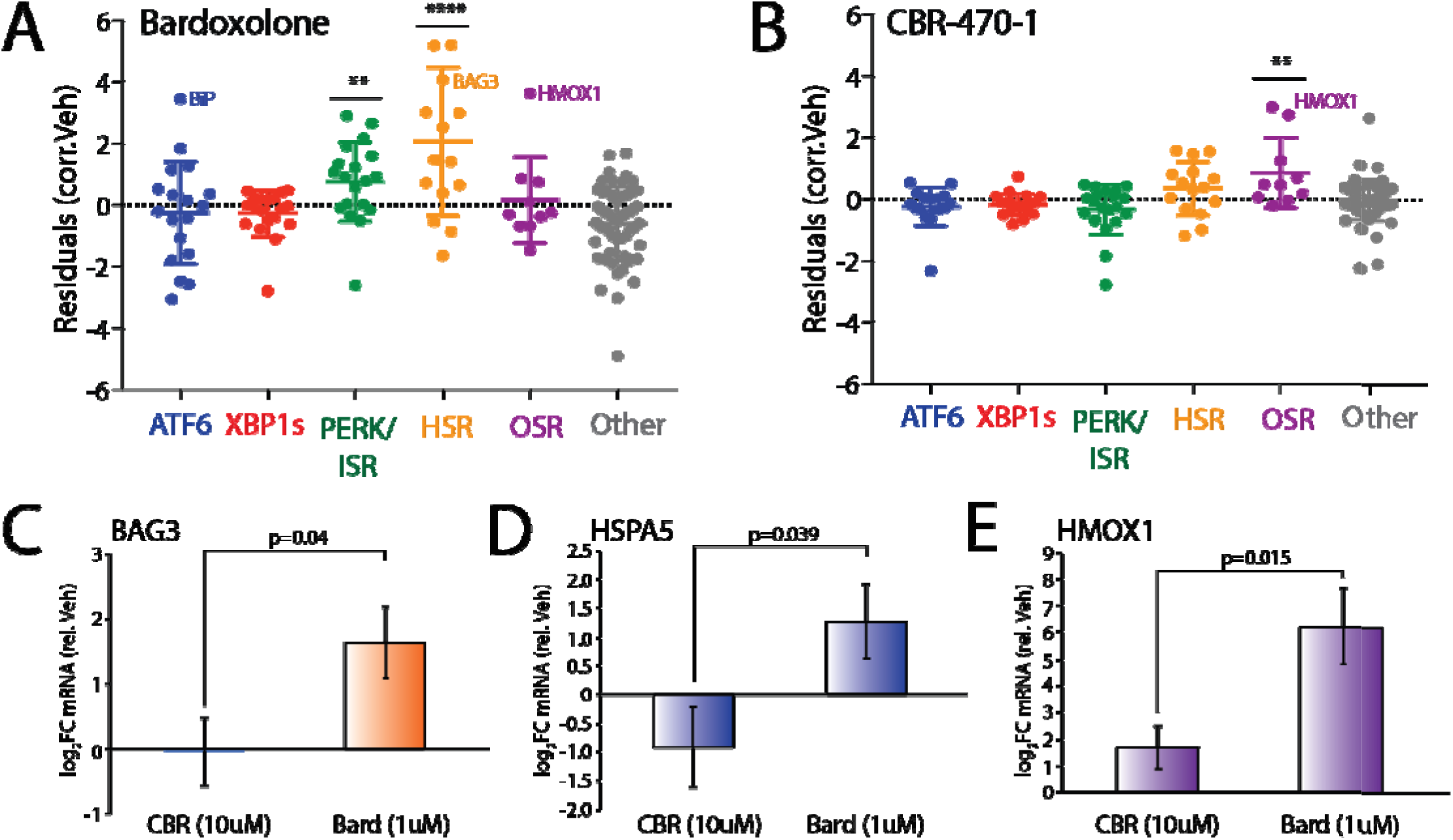
Defining the selectivity of reported NRF2 activating compounds through targeted RNAseq transcriptional profiling. A. Plot showing residuals calculated by comparing the expression of our stress-responsive gene panel between HEK293T cells treated with bardoxolone (1 μM, 24 h) or vehicle. Calculation of residuals was performed as described in Fig. 2A. Genes are grouped by target stress-responsive signaling pathway. Statistics were calculated using one-way ANOVA, Significance shown reflects comparison to “Other” target transcript set. **p<0.01, ****p<0.0001. See **Table S3** for full ANOVA table.
B. Plot showing residuals calculated by comparing the expression of our stress-responsive gene panel between HEK293T cells treated with CBR-470-1 (10 μM, 24 h) or vehicle. Calculation of residuals was performed as described in Fig. 2A. Genes are grouped by target stress-responsive signaling pathway. Statistics were calculated using one-way ANOVA, Significance shown reflects comparison to “Other” target transcript set. **p<0.01. See **Table S3** for full ANOVA table.
C. Graph showing qPCR analysis of the HSR target gene *BAG3* in HEK293T cells treated for 24 h with bardoxolone (1 μM) or CBR-470-1 (10 μM). Error bars show SEM for n= 3. P-values calculated using one-tailed Student’s t-test.
D. Graph showing qPCR analysis of the UPR (ATF6) target gene *BiP* in HEK293T cells treated for 24 h with bardoxolone (1 μM) or CBR-470-1 (10 μM). Error bars show SEM for n= 3. P-values calculated using one-tailed Student’s t-test.
E. Graph showing qPCR analysis of the OSR target gene *HMOX1* in HEK293T cells treated for 24 h with bardoxolone (1 μM) or CBR-470-1 (10 μM). Error bars show SEM for n= 3. P-values calculated using one-tailed Student’s t-test.

### Identification of selective HSR activating compounds by targeted RNAseq

Previous high-throughput screening identified numerous compounds, including compounds A3, C1, D1, and F1 (Fig. S6A), that activate a cell based reporter of the HSR-associated transcription factor HSF1 in HeLa cells ^20^. However, the selectivity of these compounds for the HSR remains to be fully defined. We used our targeted RNAseq assay to define the selectivity for these putative HSF1 activating compounds for specific HSR activation. Our results show that compound A3 strongly induced the HSR geneset (Fig. 6A) to a level comparable to that observed with dox-dependent cHSF1 activation (Fig. 2B). Compounds C1, D1, and F1 also significantly induced the HSR geneset, albeit to a lower extent (Fig. 6B-D). Interestingly, administration of these compounds also induced expression of other stress-responsive genes. This was most evident with A3, which showed robust activation of select ISR and OSR target genes such as *ATF3* and *HMOX1*, respectively, without global activation of these pathways (Fig. 6A). Similar results were observed for the other three compounds to lesser extents (Fig. 6B-D). Interestingly, both *ATF3* and *HMOX1* have been shown to be transcriptionally induced following stress-independent activation of the HSR-associated transcription factor HSF1^17^, suggesting that their increased expression in response compound treatment could, in part, reflect HSF1 activity.

**Figure 6.**
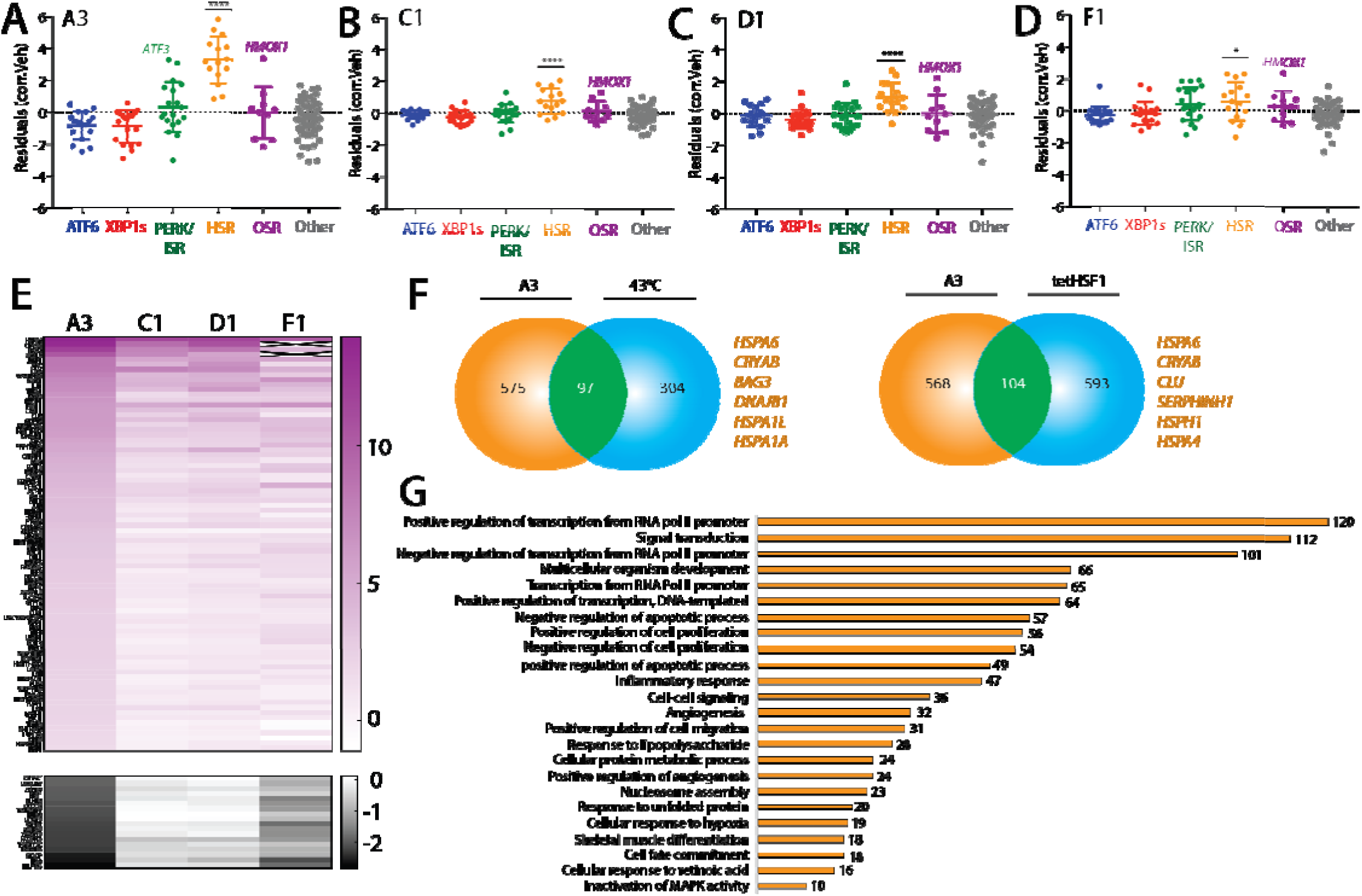
Defining the selectivity for HSF1 activating compounds identified through reporter based HTS. A. Plot showing residuals calculated by comparing the expression of our stress-responsive gene panel between HEK293T cells treated with compound A3 (10 μM, 4 h) or vehicle. Calculation of residuals was performed as described in Fig. 2A. Genes are grouped by target stress-responsive signaling pathway. Statistics were calculated using one-way ANOVA. Significance shown reflects comparison to “Other” target transcript set. ****p<0.0001. See **Table S3** for full ANOVA table.
B. Plot showing residuals calculated by comparing the expression of our stress-responsive gene panel between HEK293T cells treated with compound C1 (10 μM, 4 h) or vehicle. Calculation of residuals was performed as described in Fig. 2A. Genes are grouped by target stress-responsive signaling pathway. Statistics were calculated using one-way ANOVA. Significance shown reflects comparison to “Other” target transcript set. ****p<0.0001. See **Table S3** for full ANOVA table.
C. Plot showing residuals calculated by comparing the expression of our stress-responsive gene panel between HEK293T cells treated with compound D1 (10 μM, 4 h) or vehicle. Calculation of residuals was performed as described in Fig. 2A. Genes are grouped by target stress-responsive signaling pathway. Statistics were calculated using one-way ANOVA. Significance shown reflects comparison to “Other” target transcript set. ****p<0.0001. See **Table S3** for full ANOVA table.
D. Plot showing residuals calculated by comparing the expression of our stress-responsive gene panel between HEK293T cells treated with compound F1 (10 μM, 4 h) or vehicle. Calculation of residuals was performed as described in Fig. 2A. Genes are grouped by target stress-responsive signaling pathway. Statistics were calculated using one-way ANOVA. Significance shown reflects comparison to “Other” target transcript set. *p<0.05. See **Table S3** for full ANOVA table.
E. Heat map of fold-change transcript levels from whole-transcriptome RNAseq (relative to vehicle treated cells) for the 100 genes most significantly altered by A3 treatment in HEK293T cells (82 positive regulation, top; 10 negative regulation, bottom). The changes in these genes are also shown for HEK293T cells treated with C1, D1, and F4 (10uM for 4 hours).
F. Venn diagrams showing overlap of upregulated genes from whole-transcriptome RNAseq of HEK293T cells treated with A3 (10μM 4 hours) and HeLa cells under heat shock (left)^6^, or dox-inducible cHSF1 (right) in HEK293^trex^ cells^17^. Select genes identified in the overlap are highlighted.
G. Numbers of genes from GO analysis of significantly altered transcripts in HEK293T cells treated with compound A3 (10 μM, 4 h) relative to vehicle-treated cells. GO analysis was performed using David^59^. Entire GO analysis is reported in **Supplementary Table S5**.

To further define the selectivity of these HSR activating compounds for the HSR proteostasis transcriptional program in HEK293T cells, we performed whole transcriptome RNAseq (**Table S4**). Analysis of the top 100 most significantly altered transcripts in this whole transcriptome RNAseq data demonstrated that compound A3 induced the greatest effects on gene expression, consistent with our targeted RNAseq results (Fig. 6E). Furthermore, performing the same correlation-based geneset analysis used for targeted RNAseq revealed an identical preferential activation of the HSR in this whole transcriptome dataset (**Fig. S6B-I**). Interestingly, comparing genes induced by A3 with those induced by a 43°C heat shock^6^ or dox-dependent cHSF1^17^ activation demonstrated an overlap of ~100 genes (Fig. 6F), including many classical HSR proteostasis target genes such as *HSPA1A, DNAJB1*, and *CryAB* (**Table S4**). While this supports an A3-dependent induction of the HSR, there are a large number of genes upregulated in the whole-transcriptome data that are not upregulated by these other HSR-activating insults. GO analysis reveals that, apart from proteostasis factors, the majority of targets induced by treatment with A3 are involved in RNA polymerase II-dependent transcription (Fig. 6G). This finding is consistent with recent studies indicating that apart from direct transcriptional upregulation, HSF1 may recruit factors that increase the rate of Pol II release from its paused state in transcript promoter regions^6^. Thus, the altered expression of Pol II regulatory factors suggests that A3 could influence HSR activity by targeting RNA Pol II pause-release. While the impact of A3 on RNA Pol II could limit the further development and usage of this compound as a chemical tool to define HSR-dependent regulation of cellular proteostasis, these results demonstrate the potential for our targeted RNAseq assay to define the selectivity of prioritized compounds identified through reporter based HTS for activating specific stress-responsive proteostasis pathways.

## CONCLUDING REMARKS

The establishment of highly selective activators of stress-responsive signaling pathways provides unique opportunities to identify new roles for these pathways in regulating cellular physiology and defining their potential for correcting pathologic defects associated with diverse diseases. Here, we establish a medium-throughput targeted RNAseq assay that reports on the activation of predominant stress-responsive proteostasis pathways such as the HSR, OSR, ISR, and UPR. We demonstrate the potential for this approach to deconvolute the complex integration of stress-responsive signaling pathways in cells treated with chemical genetic or pharmacologic perturbations. Furthermore, we show that this approach is suitable for defining the selectivity of putative activators of different stress-responsive signaling pathways. These results demonstrate that this assay provides new opportunities to improve screening efforts focused on establishing pharmacologic activators of stress-responsive signaling pathways by identifying compounds or classes with high selectivity earlier in the screening pipeline (i.e., after reporter based HTS). Furthermore, this assay can be paired with medicinal chemistry to establish next generation compounds with improved selectivity and/or potency through monitoring activation of specific pathway reporter genesets. While we specifically focus on four primary stress-responsive proteostasis pathways (see Fig. 1A), this assay can be easily adapted to include new or improved genesets reporting on other signaling pathways, further improving the ability for this approach to define compound selectivity at early stages of compound development. Similarly, this assay can be adapted to other organisms, providing a unique opportunity to monitor activation of stress pathways in different tissues and define compound pharmacodynamics. Ultimately, the targeted RNAseq assay and approach described herein will improve the development of pharmacologic activators of stress-responsive signaling.

## EXPERIMENTAL PROCEDURES

### Materials and Reagents

Thapsigargin was purchased through Fisher Scientific (#50-464-295), tunicamycin was purchased through Cayman Chemical (#11089-65-9), oligomycin A was purchased from Sigma Aldrich (#75351), and paraquat was purchased from Sigma Aldrich (#36541). HSF1 and NRF2 activating compounds were generously provided by Rick Morimoto (A3, C1,D1,F1), and Michael Bollong (CBR-470-1, bardoxolone). qPCR primers used include: Bag3 (3’-TGGGAGATCAAGATCGACCC-5’, 5’-GGGCCATTGGCAGAGGATG-3’), Hspa5 (3’-GCCTGTATTTCTAGACCTGCC-5’, 5’-TTCATCTTGCCAGCCAGTTG-3’), and Hmox1 (3’-GAGTGTAAGGACCCATCGGA-5’, 5’-GCCAGCAACAAAGTGCAAG-3’) as well as RiboPro control (3’-CGTCGCCTCCTACCTGCT-5’, 5’-CCATTCAGCTCACTGATAACCTTG-3’).

### Cell lines and treatments

Stable cell lines expressing inducible cHSF1, ATF6, and XBP1s were used as previously described to activate cHSF1, ATF6, and XBP1s transcription factors^17, 34^. Cells types as listed in Table 2 were cultured in DMEM with 10% FBS, pen/strep, and glutamine at 37C, 5% CO2 in Corning 96-well tissue culture plates. Cells were treated for the indicated durations (Supplementary Table S2) with either chemical genetic activators or pharmacologics solubilized in DMSO, treatments were performed in triplicate.

### RNA extraction

RNA was extracted from HEK293T, HEK293^Dax^, and HEK293^trex^ using the Zymo Research ZR-96 Quick-RNA isolation kit as per manufacturer’s instructions. Briefly, cells were rinsed with DMSO prior to lysis, samples were then cleared of particulate matter through centrifugation. The supernatant was then subject to standard column purification steps including an on-column DNase treatment, prior to RNA elution in DNase/RNase-Free water.

### Targeted RNA-seq library preparation and sequencing

Stress-responsive sequencing libraries were prepared using the Illumna TruSeq Targeted RNA Expression technology, with targeted probes selected to create a custom gene panel to report on stress-responsive transcripts across several signaling pathways. Briefly, 50ng intact total RNA was reverse transcribed into cDNA using ProtoScript II Reverse Transcriptase (25°C 5 min, 42°C 15 min, 95°C 10 min, 4°C hold) and the targeted oligo pool was hybridized in each sample (70°C 5 min, 68°C 1 min, 65°C 2.5 min, 60°C 2.5 min, 55°C 4 min, 50°C 4 min, 45°C 4 min, 40°C 4 min, 35°C 4 min, 30°C 4 min, hold at 30°C). Hybridized products were washed using magnetic beads, extended, and amplified using i7 and i5 adapters (95°C 2 min, 28x: 98°C 30 sec, 62°C 30 sec, 72°C 60 sec, followed by 72°C for 5 minutes, 10°C hold). Libraries were cleaned and pooled prior to being loaded on the MiSeq desktop sequencer.

### Alignment and Expression Analysis

Alignment of targeted RNA-seq reads was performed using the Illumina TruSeq Targeted RNA-seq software using the custom target manifest containing sequences of targeted region sequences. Alignment of whole-transcriptome RNA-seq data was done using DNAstar Lasergene SeqManPro to the GRCh37.p13 human genome reference assembly. Aligned counts from Targeted RNAseq were median normalized and Log2 transformed prior to correlation analysis.

### Whole-transcriptome RNA-seq

HEK293T cells were treated with 10uM A3, C1, D1, or F1 for 6 hours prior to RNA isolation using the ZymoPure RNA-mini kit as per manufacturer’s instructions, including on-column DNase I treatment to remove contaminating genomic DNA. RNA was quantified by NanoDrop. Conventional RNA-seq was conducted via BGI Americas on the BGI Proprietary platform, providing single-end 50bp reads at 20 million reads per sample. Each condition was performed in triplicate.

### Gene expression correlation network graph and hierarchical clustering

Raw counts data from the triplicate RNAseq experiments for each condition were averaged and Pearson correlation coefficients were calculated for each pair of genes. We created the gene expression correlation graph by representing each gene as a vertex and connecting the vertices for the genes that had correlation coefficients ≥ 0.6. There were a few genes whose expression levels did not correlate with those of any other genes at this level. These genes were connected only to the gene with which they had the highest correlation coefficient to ensure that the network graph was fully connected. The hierarchical clustering of genes by expression pattern shown in Figure 1 was performed using the Euclidean distance between each genes’ expression level correlation coefficients with all other genes as the distance metric and single-linkage clustering as the linkage criterion. Thus, two genes that have similar sets of correlation coefficients with all other genes were most likely to cluster together. The network graph and dendrogram in Figure 1 were produced using Mathematica 11.3.

### Statistical analysis

Statistical significance of residual values from targeted RNAseq correlation analysis were calculated using standard one-way ANOVA, with the lowest p-values presented. Full ANOVA tables from our targeted RNAseq assay are included in **Table S3**. qPCR data was analyzed using one-tailed Student’s t-test.

## Supporting information

## ACKNOWLEDGEMENTS

We thank Steven Head, John Shimashita, Jeffery W. Kelly and Jessica Rosarda at The Scripps Research Institute for fruitful discussions focused on the work described in this manuscript. This work was supported by NIH AG046495 (RLW), NIH DK106582 (ETP), NIH GM101644 (ETP) and a Career Development Program Fellow Grant from the Leukemia and Lymphoma Society (Grant 5439-16; LP).

## REFERENCES

1. Hetz, C.; Mollereau, B., Disturbance of endoplasmic reticulum proteostasis in neurodegenerative diseases. Nat Rev Neurosci 2014, 15 (4), 233–49.

2. Henning, R. H.; Brundel, B., Proteostasis in cardiac health and disease. Nat Rev Cardiol 2017, 14 (11), 637–653.

3. James, H. A.; O’Neill, B. T.; Nair, K. S., Insulin Regulation of Proteostasis and Clinical Implications. Cell Metab 2017, 26 (2), 310–323.

4. Wang, S.; Kaufman, R. J., The impact of the unfolded protein response on human disease. J Cell Biol 2012, 197 (7), 857–67.

5. Wang, M.; Kaufman, R. J., Protein misfolding in the endoplasmic reticulum as a conduit to human disease. Nature 2016, 529 (7586), 326–35.

6. Mahat, D. B.; Salamanca, H. H.; Duarte, F. M.; Danko, C. G.; Lis, J. T., Mammalian Heat Shock Response and Mechanisms Underlying Its Genome-wide Transcriptional Regulation. Mol Cell 2016, 62 (1), 63–78.

7. Li, J.; Labbadia, J.; Morimoto, R. I., Rethinking HSF1 in Stress, Development, and Organismal Health. Trends Cell Biol 2017, 27 (12), 895–905.

8. Gomez-Pastor, R.; Burchfiel, E. T.; Thiele, D. J., Regulation of heat shock transcription factors and their roles in physiology and disease. Nat Rev Mol Cell Biol 2018, 19 (1), 4–19.

9. Gorrini, C.; Harris, I. S.; Mak, T. W., Modulation of oxidative stress as an anticancer strategy. Nat Rev Drug Discov 2013, 12 (12), 931–47.

10. Taguchi, K.; Motohashi, H.; Yamamoto, M., Molecular mechanisms of the Keap1-Nrf2 pathway in stress response and cancer evolution. Genes Cells 2011, 16 (2), 123–40.

11. Alam, J.; Stewart, D.; Touchard, C.; Boinapally, S.; Choi, A. M.; Cook, J. L., Nrf2, a Cap’n’Collar transcription factor, regulates induction of the heme oxygenase-1 gene. J Biol Chem 1999, 274 (37), 26071–8.

12. Walter, P.; Ron, D., The unfolded protein response: from stress pathway to homeostatic regulation. Science 2011, 334 (6059), 1081–6.

13. Han, J.; Kaufman, R. J., Physiological/pathological ramifications of transcription factors in the unfolded protein response. Genes Dev 2017, 31 (14), 1417–1438.

14. Young, S. K.; Wek, R. C., Upstream Open Reading Frames Differentially Regulate Gene-specific Translation in the Integrated Stress Response. J Biol Chem 2016, 291 (33), 16927–35.

15. Donnelly, N.; Gorman, A. M.; Gupta, S.; Samali, A., The eIF2alpha kinases: their structures and functions. Cell Mol Life Sci 2013, 70 (19), 3493–511.

16. Pakos-Zebrucka, K.; Koryga, I.; Mnich, K.; Ljujic, M.; Samali, A.; Gorman, A. M., The integrated stress response. EMBO Rep 2016, 17 (10), 1374–1395.

17. Ryno, L. M.; Genereux, J. C.; Naito, T.; Morimoto, R. I.; Powers, E. T.; Shoulders, M. D.; Wiseman, R. L., Characterizing the altered cellular proteome induced by the stress-independent activation of heat shock factor 1. ACS Chem Biol 2014, 9 (6) 1273–83.

18. Lu, P. D.; Jousse, C.; Marciniak, S. J.; Zhang, Y.; Novoa, I.; Scheuner, D.; Kaufman, R. J.; Ron, D.; Harding, H. P., Cytoprotection by pre-emptive conditional phosphorylation of translation initiation factor 2. EMBO J 2004, 23 (1), 169–79.

19. Chen, J. J.; Genereux, J. C.; Wiseman, R. L., Endoplasmic reticulum quality control and systemic amyloid disease: Impacting protein stability from the inside out. IUBMB Life 2015, 67 (6), 404–13.

20. Calamini, B.; Silva, M. C.; Madoux, F.; Hutt, D. M.; Khanna, S.; Chalfant, M. A.; Saldanha, S. A.; Hodder, P.; Tait, B. D.; Garza, D.; Balch, W. E.; Morimoto, R. I., Small-molecule proteostasis regulators for protein conformational diseases. Nat Chem Biol 2011, 8 (2), 185–96.

21. Balch, W. E.; Morimoto, R. I.; Dillin, A.; Kelly, J. W., Adapting proteostasis for disease intervention. Science 2008, 319 (5865), 916–9.

22. Cuadrado, A.; Manda, G.; Hassan, A.; Alcaraz, M. J.; Barbas, C.; Daiber, A.; Ghezzi, P.; Leon, R.; Lopez, M. G.; Oliva, B.; Pajares, M.; Rojo, A. I.; Robledinos-Anton, N.; Valverde, A. M.; Guney, E.; Schmidt, H., Transcription Factor NRF2 as a Therapeutic Target for Chronic Diseases: A Systems Medicine Approach. Pharmacol Rev 2018, 70 (2), 348–383.

23. Plate, L.; Wiseman, R. L., Regulating Secretory Proteostasis through the Unfolded Protein Response: From Function to Therapy. Trends Cell Biol 2017, 27 (10), 722–737.

24. Sweeney, P.; Park, H.; Baumann, M.; Dunlop, J.; Frydman, J.; Kopito, R.; McCampbell, A.; Leblanc, G.; Venkateswaran, A.; Nurmi, A.; Hodgson, R., Protein misfolding in neurodegenerative diseases: implications and strategies. Transl Neurodegener 2017, 6, 6.

25. Plate, L.; Cooley, C. B.; Chen, J. J.; Paxman, R. J.; Gallagher, C. M.; Madoux, F.; Genereux, J. C.; Dobbs, W.; Garza, D.; Spicer, T. P.; Scampavia, L.; Brown, S. J.; Rosen, H.; Powers, E. T.; Walter, P.; Hodder, P.; Wiseman, R. L.; Kelly, J. W., Small molecule proteostasis regulators that reprogram the ER to reduce extracellular protein aggregation. Elife 2016, 5.

26. Bollong, M. J.; Lee, G.; Coukos, J. S.; Yun, H.; Zambaldo, C.; Chang, J. W.; Chin, E. N.; Ahmad, I.; Chatterjee, A. K.; Lairson, L. L.; Schultz, P. G.; Moellering, R. E., A metabolite-derived protein modification integrates glycolysis with KEAP1-NRF2 signalling. Nature 2018, 562 (7728), 600–604.

27. Flaherty, D. P.; Miller, J. R.; Garshott, D. M.; Hedrick, M.; Gosalia, P.; Li, Y.; Milewski, M.; Sugarman, E.; Vasile, S.; Salaniwal, S.; Su, Y.; Smith, L. H.; Chung, T. D.; Pinkerton, A. B.; Aube, J.; Callaghan, M. U.; Golden, J. E.; Fribley, A. M.; Kaufman, R. J., Discovery of Sulfonamidebenzamides as Selective Apoptotic CHOP Pathway Activators of the Unfolded Protein Response. ACS Med Chem Lett 2014, 5 (12), 1278–1283.

28. Kudo, T.; Kanemoto, S.; Hara, H.; Morimoto, N.; Morihara, T.; Kimura, R.; Tabira, T.; Imaizumi, K.; Takeda, M., A molecular chaperone inducer protects neurons from ER stress. Cell Death Differ 2008, 15 (2), 364–75.

29. Salehi, A. H.; Morris, S. J.; Ho, W. C.; Dickson, K. M.; Doucet, G.; Milutinovic, S.; Durkin, J.; Gillard, J. W.; Barker, P. A., AEG3482 is an antiapoptotic compound that inhibits Jun kinase activity and cell death through induced expression of heat shock protein 70. Chem Biol 2006, 13 (2), 213–23.

30. Westerheide, S. D.; Bosman, J. D.; Mbadugha, B. N.; Kawahara, T. L.; Matsumoto, G.; Kim, S.; Gu, W.; Devlin, J. P.; Silverman, R. B.; Morimoto, R. I., Celastrols as inducers of the heat shock response and cytoprotection. J Biol Chem 2004, 279 (53), 56053–60.

31. Yang, H.; Chen, D.; Cui, Q. C.; Yuan, X.; Dou, Q. P., Celastrol, a triterpene extracted from the Chinese “Thunder of God Vine,” is a potent proteasome inhibitor and suppresses human prostate cancer growth in nude mice. Cancer Res 2006, 66 (9), 4758–65.

32. Trott, A.; West, J. D.; Klaic, L.; Westerheide, S. D.; Silverman, R. B.; Morimoto, R. I.; Morano, K. A., Activation of heat shock and antioxidant responses by the natural product celastrol: transcriptional signatures of a thiol-targeted molecule. Mol Biol Cell 2008, 19 (3), 1104–12.

33. Pronk, T. E.; van der Veen, J. W.; Vandebriel, R. J.; van Loveren, H.; de Vink, E. P.; Pennings, J. L., Comparison of the molecular topologies of stress-activated transcription factors HSF1, AP-1, NRF2, and NF-kappaB in their induction kinetics of HMOX1. Biosystems 2014, 124, 75–85.

34. Shoulders, M. D.; Ryno, L. M.; Genereux, J. C.; Moresco, J. J.; Tu, P. G.; Wu, C.; Yates, J. R., 3rd; Su, A. I.; Kelly, J. W.; Wiseman, R. L., Stress-independent activation of XBP1s and/or ATF6 reveals three functionally diverse ER proteostasis environments. Cell Rep 2013, 3 (4), 1279–92.

35. Ye, C.; Ho, D. J.; Neri, M.; Yang, C.; Kulkarni, T.; Randhawa, R.; Henault, M.; Mostacci, N.; Farmer, P.; Renner, S.; Ihry, R.; Mansur, L.; Keller, C. G.; McAllister, G.; Hild, M.; Jenkins, J.; Kaykas, A., DRUG-seq for miniaturized high-throughput transcriptome profiling in drug discovery. Nat Commun 2018, 9 (1), 4307.

36. Martin, D. P.; Miya, J.; Reeser, J. W.; Roychowdhury, S., Targeted RNA Sequencing Assay to Characterize Gene Expression and Genomic Alterations. J Vis Exp 2016, (114).

37. Koltai, H.; Weingarten-Baror, C., Specificity of DNA microarray hybridization: characterization, effectors and approaches for data correction. Nucleic Acids Res 2008, 36 (7), 2395–405.

38. Holtz, Y.; Ardisson, M.; Ranwez, V.; Besnard, A.; Leroy, P.; Poux, G.; Roumet, P.; Viader, V.; Santoni, S.; David, J., Genotyping by Sequencing Using Specific Allelic Capture to Build a High-Density Genetic Map of Durum Wheat. PLoS One 2016, 11 (5), e0154609.

39. Reeser, J. W.; Martin, D.; Miya, J.; Kautto, E. A.; Lyon, E.; Zhu, E.; Wing, M. R.; Smith, A.; Reeder, M.; Samorodnitsky, E.; Parks, H.; Naik, K. R.; Gozgit, J.; Nowacki, N.; Davies, K. D.; Varella-Garcia, M.; Yu, L.; Freud, A. G.; Coleman, J.; Aisner, D. L.; Roychowdhury, S., Validation of a Targeted RNA Sequencing Assay for Kinase Fusion Detection in Solid Tumors. J Mol Diagn 2017, 19 (5), 682–696.

40. Martinez-Jacobo, L.; Ancer-Arellano, C. I.; Ortiz-Lopez, R.; Salinas-Santander, M.; Villarreal-Villarreal, C. D.; Ancer-Rodriguez, J.; Camacho-Zamora, B.; Zomosa-Signoret, V.; Medina-De la Garza, C. E.; Ocampo-Candiani, J.; Rojas-Martinez, A., Evaluation of the Expression of Genes Associated with Inflammation and Apoptosis in Androgenetic Alopecia by Targeted RNA-Seq. Skin Appendage Disord 2018, 4 (4), 268–273.

41. Hayes, J. D.; Dinkova-Kostova, A. T., The Nrf2 regulatory network provides an interface between redox and intermediary metabolism. Trends Biochem Sci 2014, 39 (4), 199–218.

42. Lee, A. H.; Iwakoshi, N. N.; Glimcher, L. H., XBP-1 regulates a subset of endoplasmic reticulum resident chaperone genes in the unfolded protein response. Mol Cell Biol 2003, 23 (21), 7448–59.

43. Yamamoto, K.; Sato, T.; Matsui, T.; Sato, M.; Okada, T.; Yoshida, H.; Harada, A.; Mori, K., Transcriptional induction of mammalian ER quality control proteins is mediated by single or combined action of ATF6alpha and XBP1. Dev Cell 2007, 13 (3), 365–76.

44. Harding, H. P.; Zhang, Y.; Zeng, H.; Novoa, I.; Lu, P. D.; Calfon, M.; Sadri, N.; Yun, C.; Popko, B.; Paules, R.; Stojdl, D. F.; Bell, J. C.; Hettmann, T.; Leiden, J. M.; Ron, D., An integrated stress response regulates amino acid metabolism and resistance to oxidative stress. Mol Cell 2003, 11 (3), 619–33.

45. Dengler, V. L.; Galbraith, M.; Espinosa, J. M., Transcriptional regulation by hypoxia inducible factors. Crit Rev Biochem Mol Biol 2014, 49 (1), 1–15.

46. Raskatov, J. A.; Meier, J. L.; Puckett, J. W.; Yang, F.; Ramakrishnan, P.; Dervan, P. B., Modulation of NF-kappaB-dependent gene transcription using programmable DNA minor groove binders. Proc Natl Acad Sci U S A 2012, 109 (4), 1023–8.

47. Zhao, Q.; Wang, J.; Levichkin, I. V.; Stasinopoulos, S.; Ryan, M. T.; Hoogenraad, N. J., A mitochondrial specific stress response in mammalian cells. EMBO J 2002, 21 (17), 4411–9.

48. Aldridge, J. E.; Horibe, T.; Hoogenraad, N. J., Discovery of genes activated by the mitochondrial unfolded protein response (mtUPR) and cognate promoter elements. PLoS One 2007, 2 (9), e874.

49. Melber, A.; Haynes, C. M., UPR(mt) regulation and output: a stress response mediated by mitochondrial-nuclear communication. Cell Res 2018, 28 (3), 281–295.

50. Kim, J. M.; Kim, H. G.; Son, C. G., Tissue-Specific Profiling of Oxidative Stress-Associated Transcriptome in a Healthy Mouse Model. Int J Mol Sci 2018, 19 (10).

51. Miao, L.; St Clair, D. K., Regulation of superoxide dismutase genes: implications in disease. Free Radic Biol Med 2009, 47 (4), 344–56.

52. Richard, E.; Desviat, L. R.; Ugarte, M.; Perez, B., Oxidative stress and apoptosis in homocystinuria patients with genetic remethylation defects. J Cell Biochem 2013, 114 (1), 183–91.

53. Leclerc, D.; Rozen, R., Endoplasmic reticulum stress increases the expression of methylenetetrahydrofolate reductase through the IRE1 transducer. J Biol Chem 2008, 283 (6), 3151–60.

54. Druwe, I. L.; Vaillancourt, R. R., Influence of arsenate and arsenite on signal transduction pathways: an update. Arch Toxicol 2010, 84 (8), 585–96.

55. Qureshi, M. A.; Haynes, C. M.; Pellegrino, M. W., The mitochondrial unfolded protein response: Signaling from the powerhouse. J Biol Chem 2017, 292 (33), 13500–13506.

56. Chin, M. P.; Bakris, G. L.; Block, G. A.; Chertow, G. M.; Goldsberry, A.; Inker, L. A.; Heerspink, H. J. L.; O’Grady, M.; Pergola, P. E.; Wanner, C.; Warnock, D. G.; Meyer, C. J., Bardoxolone Methyl Improves Kidney Function in Patients with Chronic Kidney Disease Stage 4 and Type 2 Diabetes: Post-Hoc Analyses from Bardoxolone Methyl Evaluation in Patients with Chronic Kidney Disease and Type 2 Diabetes Study. Am J Nephrol 2018, 47 (1), 40–47.

57. Yore, M. M.; Kettenbach, A. N.; Sporn, M. B.; Gerber, S. A.; Liby, K. T., Proteomic analysis shows synthetic oleanane triterpenoid binds to mTOR. PLoS One 2011, 6 (7), e22862.

58. Gibellini, L.; Pinti, M.; Bartolomeo, R.; De Biasi, S.; Cormio, A.; Musicco, C.; Carnevale, G.; Pecorini, S.; Nasi, M.; De Pol, A.; Cossarizza, A., Inhibition of Lon protease by triterpenoids alters mitochondria and is associated to cell death in human cancer cells. Oncotarget 2015, 6 (28), 25466–83.

59. Huang, D. W.; Sherman, B. T.; Tan, Q.; Collins, J. R.; Alvord, W. G.; Roayaei, J.; Stephens, R.; Baseler, M. W.; Lane, H. C.; Lempicki, R. A., The DAVID Gene Functional Classification Tool: a novel biological module-centric algorithm to functionally analyze large gene lists. Genome Biol 2007, 8 (9), R183.

