## Supplementary material for "Deconvoluting Stress-Responsive Proteostasis Signaling Pathways for Pharmacologic Activation using Targeted RNA-sequencing"

Running Title: Targeted RNAseq assay for stress-signaling

Keywords: Targeted RNAseq; unfolded protein response; heat shock response; HSF1; oxidative stress response; small molecule activator; screening

#### SUPPLEMENTAL FIGURE LEGENDS

##### Figure S1 (Supplement to Fig. 1)

- **A.** Percentage of total aligned reads reporting on the selected 150 target genes of interest for Targeted RNAseq, or whole-transcriptome RNAseq from HEK293$^{TREX}$ cells expressing dox-inducible cHSF1[8], or HEK293$^{DAX}$ cells expressing TMP-inducible ATF6 and dox-inducible XBP1s[15]
- **B.** Average reads per individual gene in our Targeted RNAseq assay across all treatment conditions.
- **C.** Total reads per individual gene in our Targeted RNAseq assay across all treatment conditions.
- **D.** Example correlation analysis from our Targted RNAseq assay showing the log2 normalized aligned counts from two replicates of DMSO-treated HEK293T cells. The tight correlation demonstrates the reproducibility of this assay across individual replicates.
- **E.** Average $R^2$ values from correlations of three technical replicates (calculated as in **Fig. S1D**) for each treatment condition used in our Targted RNAseq assay (see **Table 2** and **Table S1**).

##### Figure S2 (Supplement to Fig. 2)

- **A.** $Log_2$ normalized aligned transcript counts for HEK293$^{TREX}$ cells expressing doxycycline (dox)-inducible cHSF1 treated with 2.25 μM dox (y-axis) or vehicle (x-axis) for 16 h. Aligned transcript counts represent averages from three independent replicates quantified from published whole transcriptome RNAseq {Ryno, 2014 #73}. All identified genes are HSR target genes.
- **B.** Plot showing residuals calculated by comparing the expression of our panel of stress-responsive genes between HEK293$^{TREX}$ cells expressing dox-inducible cHSF1 following 16 h treatment with dox (2.25 μM) or vehicle. Calculation of residuals was performed as described in **Fig. 2A**. Statistics were calculated using one-way ANOVA. Significance shown reflects comparison to “Other” target transcript set. ****p<0.0001. See **Table S3** for full ANOVA table.
- **C.** Residual values per target gene from whole-transcriptome RNAseq data (x-axis){Ryno, 2014 #73} vs. Targeted RNAseq (y-axis) in HEK293$^{TREX}$ cells expressing dox-inducible cHSF1 following 16 h treatment with dox (2.25μM doxycycline).

##### Figure S3 (Supplement to Fig. 3)

**A.** $Log_2$ normalized aligned transcript counts for $HEK293^{DAX}$ cells treated with 1 μM Thapsigargin (y-axis) or vehicle (x-axis) for 4 h. Aligned transcript counts represent averages from three independent replicates quantified from our targeted RNAseq data. All identified genes are UPR target genes.

**B.** Plot showing residuals calculated by comparing the expression of our panel of stress-responsive genes between $HEK293^{DAX}$ cells following 4 h treatment with Tg (1 μM; induces UPR) and vehicle. Calculation of residuals was performed as described in **Fig. 2A**. Statistics were calculated using one-way ANOVA. Significance shown reflects comparison to "Other" target transcript set. ****p<0.0001. See **Table S3** for full ANOVA table.

**C.** Residual values per target gene from whole-transcriptome RNAseq data (x-axis){Shoulders, 2013 #7} vs. Targeted RNAseq (y-axis) for $HEK293^{DAX}$ cells following treatment with trimethoprim (10 μM, 4 h; activates DHFR.ATF6).

**D.** Residual values per target gene from whole-transcriptome RNAseq data (x-axis){Shoulders, 2013 #7} vs. Targeted RNAseq (y-axis) for $HEK293^{DAX}$ cells following treatment with doxycycline (1μg/mL μM, 4 h; activates dox-inducible XBP1s).

**E.** Residual values per target gene from whole-transcriptome RNAseq data (x-axis){Shoulders, 2013 #7} vs. Targeted RNAseq (y-axis) for $HEK293^{DAX}$ cells following treatment with both trimethoprim (10 μM, 4 h; activates DHFR.ATF6) and doxycycline (1μg/mL μM, 4 h; activates dox-inducible XBP1s).

**Figure S4 (Supplement to Fig. 4)**

Plot showing residuals calculated by comparing the expression of our stress-responsive gene panel between HEK293T cells following treatment with paraquat (PQ; 400 μM, 24 h) or vehicle. Calculation of residuals was performed as described in **Fig. 2A**. Genes are grouped by target stress-responsive signaling pathway. Statistics were calculated using one-way ANOVA, significance shown reflects comparison to "Other" target transcript set. See **Table S3** for full ANOVA table.

**Figure S5 (Supplement to Fig 5).**

**A.** Structures of the putative NRF2 activating compounds bardoxolone and CBR-470-1.

B. Graph showing $log_2$ fold change normalized counts of the HSR target gene *BAG3* in HEK293T cells treated with bardoxolone (1 μM; 24 h) or CBR-470-1 (10 μM; 24 h), as calculated from our targeted RNAseq assay. Error bars show SEM for n=3 independent experiments. P-values calculated using one-tailed Student's t-test.

C. Graph showing $log_2$ fold change normalized counts of the UPR (ATF6) target gene *BIP* in HEK293T cells treated with bardoxolone (1 μM; 24 h) or CBR-470-1 (10 μM; 24 h), as calculated from our targeted RNAseq assay. Error bars show SEM for n=3 independent experiments. P-values calculated using one-tailed Student's t-test.

D. Graph showing $log_2$ fold change normalized counts of the OSR target gene *HMOX1* in HEK293T cells treated with bardoxolone (1 μM; 24 h) or CBR-470-1 (10 μM; 24 h), as calculated from our targeted RNAseq assay. Error bars show SEM for n=3 independent experiments. P-values calculated using one-tailed Student's t-test.

**Figure S6 (Supplement to Fig. 6)**

A. Structures of the four putative HSF1 activating compounds A3, C1, D1, and F1.

B. Plot showing residuals calculated by comparing the expression of our stress-responsive gene panel between HEK293T cells treated with compound A3 (10 μM, 4 h) or vehicle from whole-transcriptome RNAseq. Calculation of residuals was performed as described in **Fig. 2A**. Genes are grouped by target stress-responsive signaling pathway. Statistics were calculated using one-way ANOVA. Significance shown reflects comparison to "Other" target transcript set. ****p<0.0001.

C. Plot showing residuals calculated by comparing the expression of our stress-responsive gene panel between HEK293T cells treated with compound C1 (10 μM, 4 h) or vehicle from whole-transcriptome RNAseq. Calculation of residuals was performed as described in **Fig. 2A**. Genes are grouped by target stress-responsive signaling pathway. Statistics were calculated using one-way ANOVA. Significance shown reflects comparison to "Other" target transcript set. ****p<0.0001.

D. Plot showing residuals calculated by comparing the expression of our stress-responsive gene panel between HEK293T cells treated with compound D1 (10 μM, 4 h) or vehicle from whole-transcriptome RNAseq. Calculation of residuals was performed as described in **Fig. 2A**. Genes are grouped by target stress-responsive signaling pathway. Statistics were calculated using one-way ANOVA. Significance shown reflects comparison to “Other” target transcript set. ****p<0.0001.

**E.** Plot showing residuals calculated by comparing the expression of our stress-responsive gene panel between HEK293T cells treated with compound F1 (10 μM, 4 h) or vehicle from whole-transcriptome RNAseq. Calculation of residuals was performed as described in **Fig. 2A**. Genes are grouped by target stress-responsive signaling pathway. Statistics were calculated using one-way ANOVA. Significance shown reflects comparison to “Other” target transcript set. ****p<0.0001.

**F.** One-way ANOVA statistical analysis for grouped residual values in (**Fig. S6B**).

**G.** One-way ANOVA statistical analysis for grouped residual values in (**Fig. S6C**).

**H.** One-way ANOVA statistical analysis for grouped residual values in (**Fig. S6D**).

**I.** One-way ANOVA statistical analysis for grouped residual values in (**Fig. S6E**).

### FIGURE S1

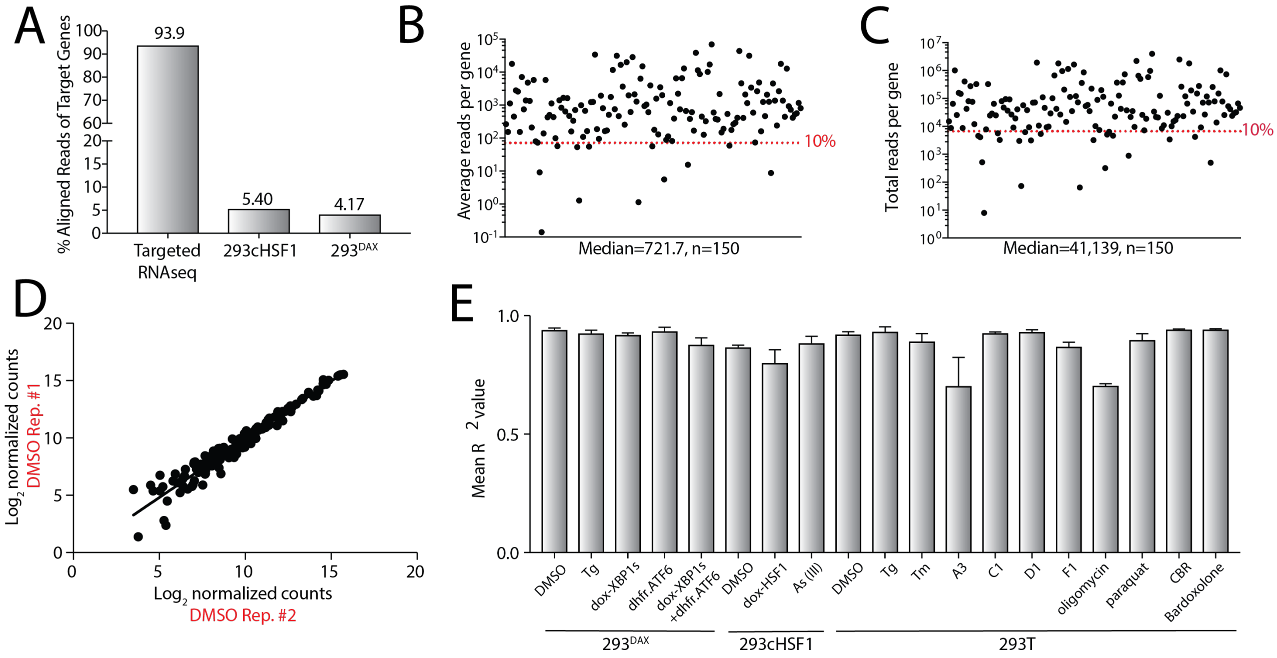

A
% Aligned Reads of Target Genes
93.9
5.40
4.17
Targeted RNAseq
293cHSF1
293DAX
B
Average reads per gene
10%
Median=721.7, n=150
C
Total reads per gene
10%
Median=41,139, n=150
D
Log2 normalized counts
DMSO Rep. #1
Log2 normalized counts
DMSO Rep. #2
E
Mean R2 value
DMSO
Tg
dox-XBP1s
dhfr.ATF6
dox-XBP1s +dhfr.ATF6
DMSO
dox-HSF1
As (III)
DMSO
Tg
Tm
A3
C1
D1
F1
oligomycin
paraquat
CBR
Bardoxolone
293DAX
293cHSF1
293T

**FIGURE S2**

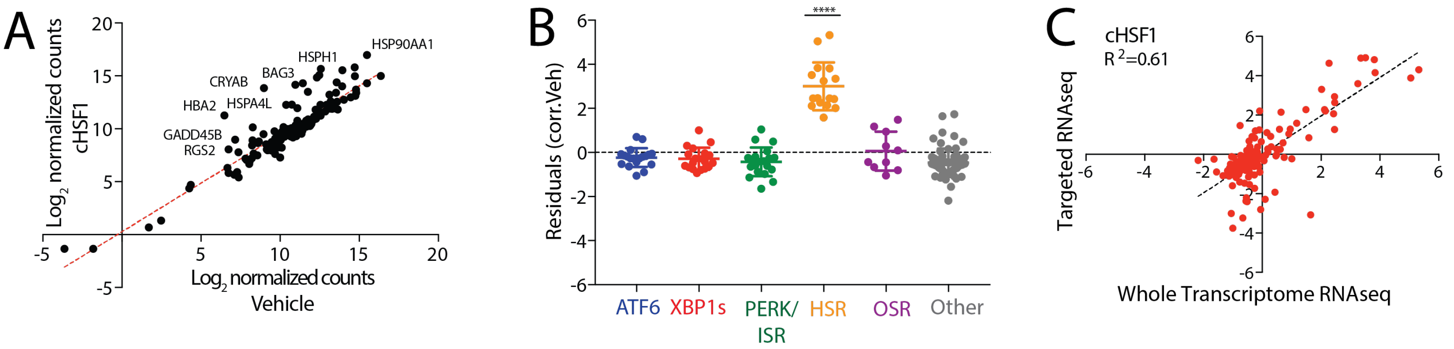
A
Log2 normalized counts
cHSF1
Log2 normalized counts
Vehicle
HSP90AA1
HSPH1
BAG3
CRYAB
HBA2
HSPA4L
GADD45B
RGS2
B
Residuals (corr.Veh)
****
ATF6
XBP1s
PERK/
ISR
HSR
OSR
Other
C
cHSF1
R 2=0.61
Targeted RNAseq
Whole Transcriptome RNAseq

**FIGURE S3**

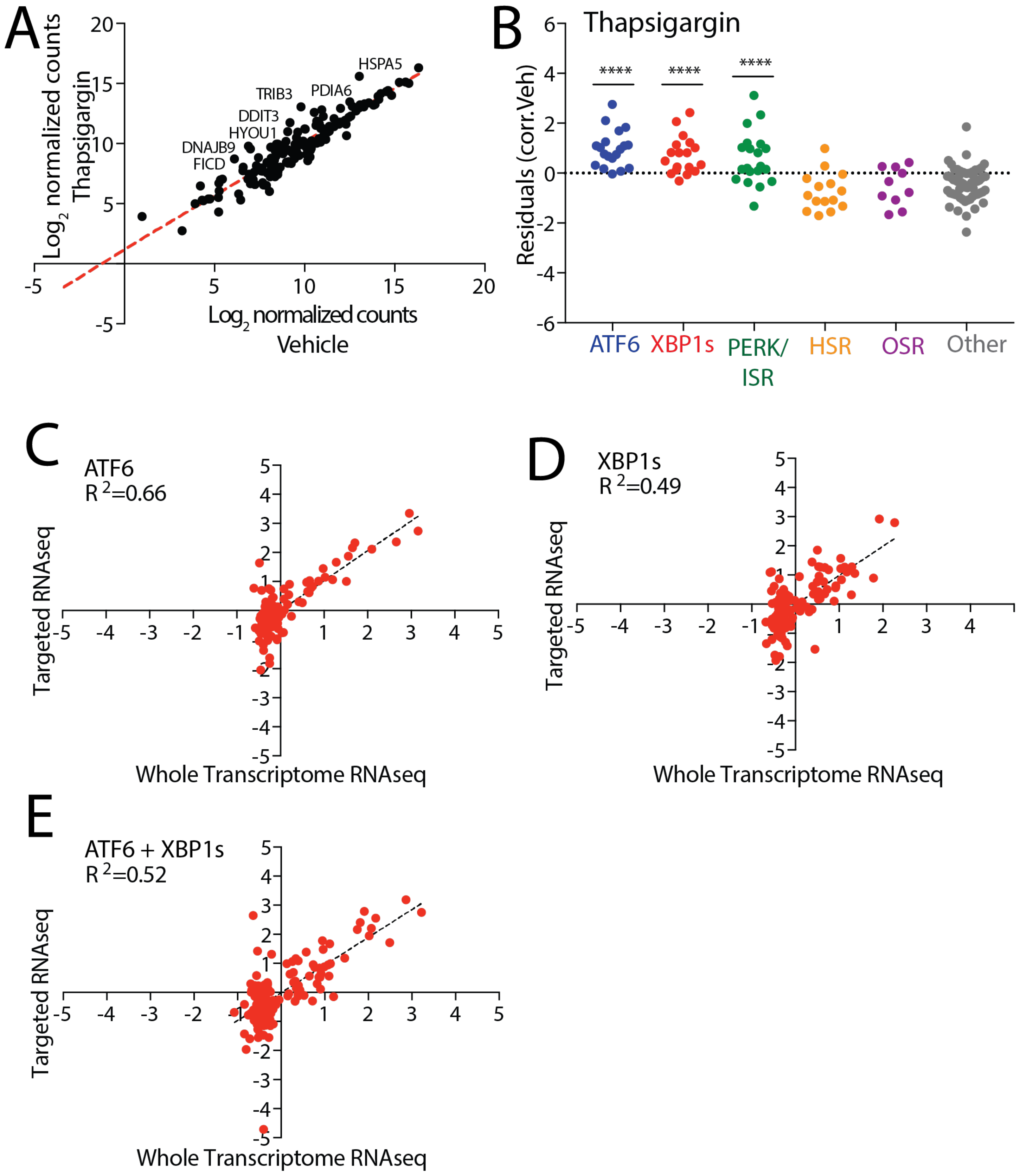

A
Log2 normalized counts
Thapsigargin
Log2 normalized counts
Vehicle
HSPA5
PDIA6
TRIB3
DDIT3
HYOU1
DNAJB9
FICD
B
Thapsigargin
Residuals (corr.Veh)
****
****
****
ATF6
XBP1s
PERK/
ISR
HSR
OSR
Other
C
ATF6
R 2=0.66
Targeted RNAseq
Whole Transcriptome RNAseq
D
XBP1s
R 2=0.49
Targeted RNAseq
Whole Transcriptome RNAseq
E
ATF6 + XBP1s
R 2=0.52
Targeted RNAseq
Whole Transcriptome RNAseq

**FIGURE S4**

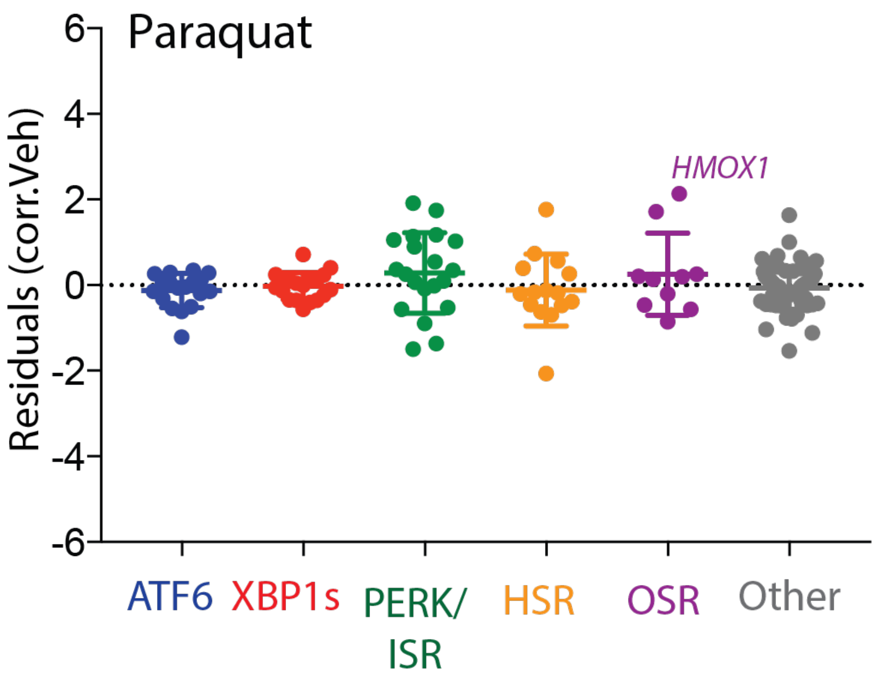

Paraquat
Residuals (corr.Veh)
6
4
2
0
-2
-4
-6
*HMOX1*
ATF6
XBP1s
PERK/ ISR
HSR
OSR
Other

**FIGURE S5**

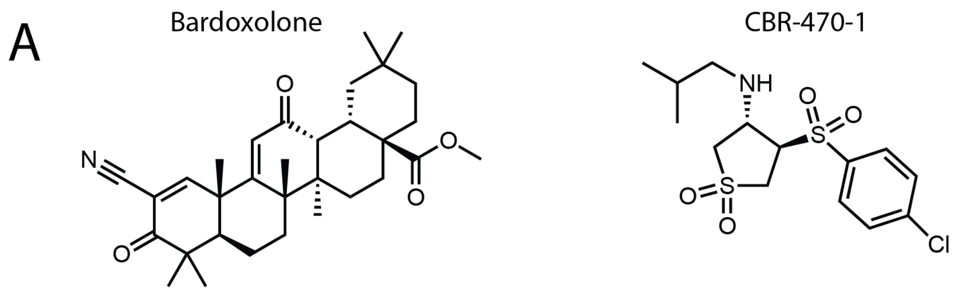
A
Bardoxolone
CBR-470-1
N
O
O
O
O
NH
O
S
O
O
S
O
Cl


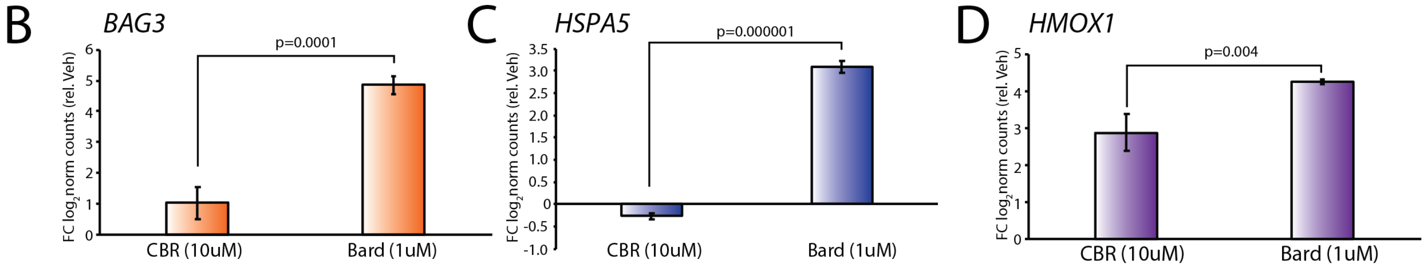
B
BAG3
p=0.0001
FC log2 norm counts (rel. Veh)
6
5
4
3
2
1
0
CBR (10uM)
Bard (1uM)
C
HSPA5
p=0.000001
FC log2 norm counts (rel. Veh)
3.5
3.0
2.5
2.0
1.5
1.0
0.5
0
-0.5
-1.0
CBR (10uM)
Bard (1uM)
D
HMOX1
p=0.004
FC log2 norm counts (rel. Veh)
5
4
3
2
1
0
CBR (10uM)
Bard (1uM)

**FIGURE S6**

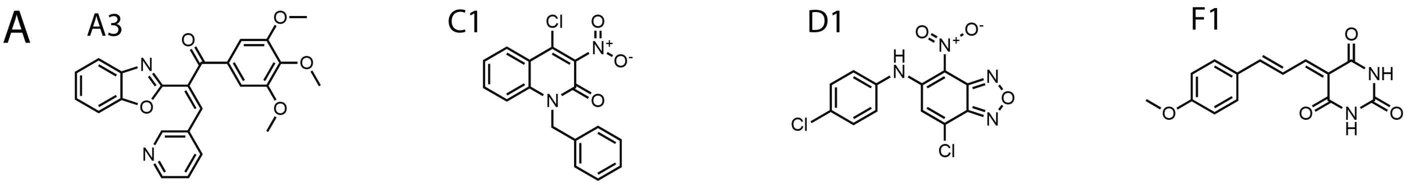


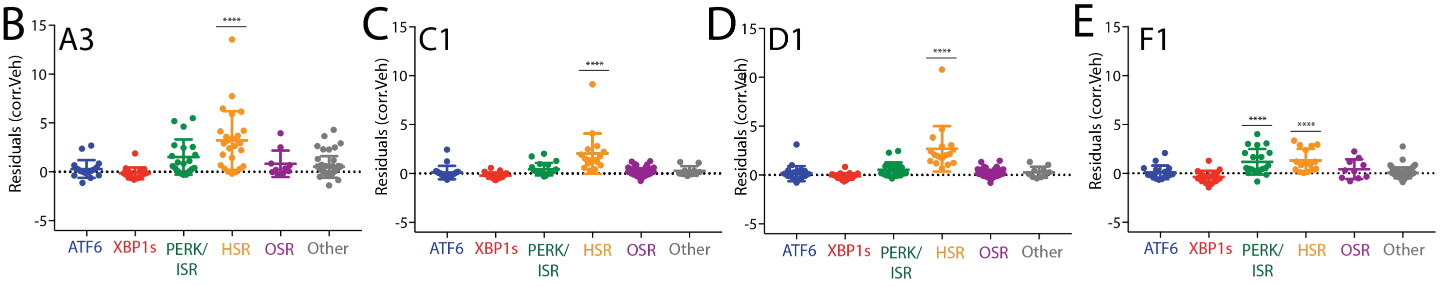


F

A3

| Comparison | Mean Diff. | Summary | Adjusted P Value |
|---|---|---|---|
| ATF6 vs. XBP1s | 0.4412 | ns | 0.9640 |
| ATF6 vs. PERK/ISR | -1.22 | ns | 0.2058 |
| ATF6 vs. HSR | -2.911 | **** | <0.0001 |
| ATF6 vs. OSR | -0.5258 | ns | 0.9656 |
| ATF6 vs. Other | -0.2103 | ns | 0.9971 |
| XBP1s vs. PERK/ISR | -1.662 | * | 0.0265 |
| XBP1s vs. HSR | -3.352 | **** | <0.0001 |
| XBP1s vs. OSR | -0.9671 | ns | 0.6731 |
| XBP1s vs. Other | -0.6515 | ns | 0.6901 |
| PERK/ISR vs. HSR | -1.691 | * | 0.0106 |
| PERK/ISR vs. OSR | 0.6945 | ns | 0.8897 |
| PERK/ISR vs. Other | 1.01 | ns | 0.1986 |
| HSR vs. OSR | 2.385 | ** | 0.0024 |
| HSR vs. Other | 2.701 | **** | <0.0001 |
| OSR vs. Other | 0.3155 | ns | 0.9940 |

G

C1

| Comparison | Mean Diff. | Summary | Adjusted P Value |
|---|---|---|---|
| ATF6 vs. XBP1s | 0.3144 | ns | 0.8591 |
| ATF6 vs. PERK/ISR | -0.3333 | ns | 0.8188 |
| ATF6 vs. HSR | -1.919 | **** | <0.0001 |
| ATF6 vs. OSR | -0.1605 | ns | 0.9965 |
| ATF6 vs. Other | -0.02284 | ns | >0.9999 |
| XBP1s vs. PERK/ISR | -0.6476 | ns | 0.1633 |
| XBP1s vs. HSR | -2.233 | **** | <0.0001 |
| XBP1s vs. OSR | -0.4749 | ns | 0.7005 |
| XBP1s vs. Other | -0.3372 | ns | 0.6514 |
| PERK/ISR vs. HSR | -1.585 | **** | <0.0001 |
| PERK/ISR vs. OSR | 0.1727 | ns | 0.9949 |
| PERK/ISR vs. Other | 0.3104 | ns | 0.7102 |
| HSR vs. OSR | 1.758 | **** | <0.0001 |
| HSR vs. Other | 1.896 | **** | <0.0001 |
| OSR vs. Other | 0.1377 | ns | 0.9968 |

H

D1

| Comparison | Mean Diff. | Summary | Adjusted P Value |
|---|---|---|---|
| ATF6 vs. XBP1s | 0.2788 | ns | 0.9329 |
| ATF6 vs. PERK/ISR | -0.3842 | ns | 0.7707 |
| ATF6 vs. HSR | -2.548 | **** | <0.0001 |
| ATF6 vs. OSR | -0.1388 | ns | 0.9988 |
| ATF6 vs. Other | 0.003854 | ns | >0.9999 |
| XBP1s vs. PERK/ISR | -0.663 | ns | 0.2419 |
| XBP1s vs. HSR | -2.827 | **** | <0.0001 |
| XBP1s vs. OSR | -0.4176 | ns | 0.8642 |
| XBP1s vs. Other | -0.275 | ns | 0.8762 |
| PERK/ISR vs. HSR | -2.164 | **** | <0.0001 |
| PERK/ISR vs. OSR | 0.2454 | ns | 0.9844 |
| PERK/ISR vs. Other | 0.388 | ns | 0.6012 |
| HSR vs. OSR | 2.409 | **** | <0.0001 |
| HSR vs. Other | 2.552 | **** | <0.0001 |
| OSR vs. Other | 0.1427 | ns | 0.9978 |

I

F1

| Comparison | Mean Diff. | Summary | Adjusted P Value |
|---|---|---|---|
| ATF6 vs. XBP1s | 0.461 | ns | 0.5437 |
| ATF6 vs. PERK/ISR | -1.077 | ** | 0.0015 |
| ATF6 vs. HSR | -1.22 | *** | 0.0005 |
| ATF6 vs. OSR | -0.3262 | ns | 0.9190 |
| ATF6 vs. Other | 0.04627 | ns | >0.9999 |
| XBP1s vs. PERK/ISR | -1.538 | **** | <0.0001 |
| XBP1s vs. HSR | -1.681 | **** | <0.0001 |
| XBP1s vs. OSR | -0.7872 | ns | 0.1556 |
| XBP1s vs. Other | -0.4147 | ns | 0.4099 |
| PERK/ISR vs. HSR | -0.1437 | ns | 0.9955 |
| PERK/ISR vs. OSR | 0.7504 | ns | 0.1893 |
| PERK/ISR vs. Other | 1.123 | **** | <0.0001 |
| HSR vs. OSR | 0.894 | ns | 0.0887 |
| HSR vs. Other | 1.266 | **** | <0.0001 |
| OSR vs. Other | 0.3724 | ns | 0.7784 |

**Table S1. Treatment conditions for Targeted RNAseq with concentrations and treatment durations** (see Excel Spreadsheet)

**Table S2 Aligned counts from Targeted RNAseq** (see Excel Spreadsheet)

**Table S3. ANOVA Statistical Analysis from Targeted RNAseq**

HSF1

| Comparison | Mean Diff. | Summary | Adjusted P Value |
|---|---|---|---|
| ATF6 vs. XBP1s | -0.2735 | ns | 0.9671 |
| ATF6 vs. PERK/ISR | -0.2218 | ns | 0.9878 |
| ATF6 vs. HSR | -3.863 | **** | <0.0001 |
| ATF6 vs. OSR | -0.421 | ns | 0.9097 |
| ATF6 vs. Other | -0.1022 | ns | 0.9991 |
| XBP1s vs. PERK/ISR | 0.05166 | ns | >0.9999 |
| XBP1s vs. HSR | -3.59 | **** | <0.0001 |
| XBP1s vs. OSR | -0.1475 | ns | 0.9992 |
| XBP1s vs. Other | 0.1712 | ns | 0.9894 |
| PERK/ISR vs. HSR | -3.641 | **** | <0.0001 |
| PERK/ISR vs. OSR | -0.1992 | ns | 0.9968 |
| PERK/ISR vs. Other | 0.1196 | ns | 0.9982 |
| HSR vs. OSR | 3.442 | **** | <0.0001 |
| HSR vs. Other | 3.761 | **** | <0.0001 |
| OSR vs. Other | 0.3187 | ns | 0.9490 |

XBP1s

| Comparison | Mean Diff. | Summary | Adjusted P Value |
|---|---|---|---|
| ATF6 vs. XBP1s | -0.5015 | ns | 0.1807 |
| ATF6 vs. PERK/ISR | 0.7097 | * | 0.0123 |
| ATF6 vs. HSR | 1.016 | *** | 0.0002 |
| ATF6 vs. OSR | 1.119 | *** | 0.0003 |
| ATF6 vs. Other | 0.8789 | **** | <0.0001 |
| XBP1s vs. PERK/ISR | 1.211 | **** | <0.0001 |
| XBP1s vs. HSR | 1.517 | **** | <0.0001 |
| XBP1s vs. OSR | 1.62 | **** | <0.0001 |
| XBP1s vs. Other | 1.38 | **** | <0.0001 |
| PERK/ISR vs. HSR | 0.3062 | ns | 0.7370 |
| PERK/ISR vs. OSR | 0.4091 | ns | 0.5805 |
| PERK/ISR vs. Other | 0.1692 | ns | 0.9128 |
| HSR vs. OSR | 0.1029 | ns | 0.9988 |
| HSR vs. Other | -0.137 | ns | 0.9775 |
| OSR vs. Other | -0.2399 | ns | 0.8867 |

ATF6

| Comparison | Mean Diff. | Summary | Adjusted P Value |
|---|---|---|---|
| ATF6 vs. XBP1s | 1.546 | **** | <0.0001 |
| ATF6 vs. PERK/ISR | 1.717 | **** | <0.0001 |
| ATF6 vs. HSR | 1.772 | **** | <0.0001 |
| ATF6 vs. OSR | 1.891 | **** | <0.0001 |
| ATF6 vs. Other | 1.737 | **** | <0.0001 |
| XBP1s vs. PERK/ISR | 0.1717 | ns | 0.9412 |
| XBP1s vs. HSR | 0.2259 | ns | 0.8722 |
| XBP1s vs. OSR | 0.3449 | ns | 0.6569 |
| XBP1s vs. Other | 0.1916 | ns | 0.8117 |
| PERK/ISR vs. HSR | 0.05413 | ns | 0.9998 |
| PERK/ISR vs. OSR | 0.1732 | ns | 0.9727 |
| PERK/ISR vs. Other | 0.01982 | ns | >0.9999 |
| HSR vs. OSR | 0.119 | ns | 0.9961 |
| HSR vs. Other | -0.03431 | ns | >0.9999 |
| OSR vs. Other | -0.1534 | ns | 0.9721 |

XBP1s+ATF6

| Comparison | Mean Diff. | Summary | Adjusted P Value |
|---|---|---|---|
| ATF6 vs. XBP1s | 0.6466 | ns | 0.1510 |
| ATF6 vs. PERK/ISR | 1.886 | **** | <0.0001 |
| ATF6 vs. HSR | 1.989 | **** | <0.0001 |
| ATF6 vs. OSR | 2.09 | **** | <0.0001 |
| ATF6 vs. Other | 1.808 | **** | <0.0001 |
| XBP1s vs. PERK/ISR | 1.239 | **** | <0.0001 |
| XBP1s vs. HSR | 1.343 | **** | <0.0001 |
| XBP1s vs. OSR | 1.444 | *** | 0.0002 |
| XBP1s vs. Other | 1.161 | **** | <0.0001 |
| PERK/ISR vs. HSR | 0.1038 | ns | 0.9991 |
| PERK/ISR vs. OSR | 0.2048 | ns | 0.9872 |
| PERK/ISR vs. Other | -0.07768 | ns | 0.9991 |
| HSR vs. OSR | 0.101 | ns | 0.9997 |
| HSR vs. Other | -0.1815 | ns | 0.9723 |
| OSR vs. Other | -0.2825 | ns | 0.9135 |

As (III)

| Comparison | Mean Diff. | Summary | Adjusted P Value |
|---|---|---|---|
| ATF6 vs. XBP1s | -0.5934 | ns | 0.7071 |
| ATF6 vs. PERK/ISR | -1.764 | *** | 0.0005 |
| ATF6 vs. HSR | -2.037 | **** | <0.0001 |
| ATF6 vs. OSR | -2.288 | *** | 0.0001 |
| ATF6 vs. Other | -0.4811 | ns | 0.7045 |
| XBP1s vs. PERK/ISR | -1.17 | ns | 0.0593 |
| XBP1s vs. HSR | -1.443 | * | 0.0138 |
| XBP1s vs. OSR | -1.695 | * | 0.0112 |
| XBP1s vs. Other | 0.1123 | ns | 0.9994 |
| PERK/ISR vs. HSR | -0.2728 | ns | 0.9886 |
| PERK/ISR vs. OSR | -0.5245 | ns | 0.8993 |
| PERK/ISR vs. Other | 1.283 | ** | 0.0026 |
| HSR vs. OSR | -0.2517 | ns | 0.9965 |
| HSR vs. Other | 1.556 | *** | 0.0004 |
| OSR vs. Other | 1.807 | *** | 0.0008 |

Oligomycin

| Comparison | Mean Diff. | Summary | Adjusted P Value |
|---|---|---|---|
| ATF6 vs. XBP1s | -0.858 | ns | 0.3063 |
| ATF6 vs. PERK/ISR | -1.496 | ** | 0.0055 |
| ATF6 vs. HSR | 0.6632 | ns | 0.6571 |
| ATF6 vs. OSR | -0.633 | ns | 0.7940 |
| ATF6 vs. Other | -0.1927 | ns | 0.9928 |
| XBP1s vs. PERK/ISR | -0.6376 | ns | 0.6083 |
| XBP1s vs. HSR | 1.521 | ** | 0.0067 |
| XBP1s vs. OSR | 0.225 | ns | 0.9972 |
| XBP1s vs. Other | 0.6653 | ns | 0.3185 |
| PERK/ISR vs. HSR | 2.159 | **** | <0.0001 |
| PERK/ISR vs. OSR | 0.8626 | ns | 0.4800 |
| PERK/ISR vs. Other | 1.303 | ** | 0.0013 |
| HSR vs. OSR | -1.296 | ns | 0.1134 |
| HSR vs. Other | -0.8559 | ns | 0.1614 |
| OSR vs. Other | 0.4403 | ns | 0.9019 |

Paraquat

| Comparison | Mean Diff. | Summary | Adjusted P Value |
|---|---|---|---|
| ATF6 vs. XBP1s | -0.1011 | ns | 0.9965 |
| ATF6 vs. PERK/ISR | -0.4086 | ns | 0.3509 |
| ATF6 vs. HSR | -0.01337 | ns | >0.9999 |
| ATF6 vs. OSR | -0.3808 | ns | 0.6450 |
| ATF6 vs. Other | -0.06267 | ns | 0.9991 |
| XBP1s vs. PERK/ISR | -0.3075 | ns | 0.6508 |
| XBP1s vs. HSR | 0.08775 | ns | 0.9986 |
| XBP1s vs. OSR | -0.2797 | ns | 0.8659 |
| XBP1s vs. Other | 0.03845 | ns | >0.9999 |
| PERK/ISR vs. HSR | 0.3952 | ns | 0.4470 |
| PERK/ISR vs. OSR | 0.02784 | ns | >0.9999 |
| PERK/ISR vs. Other | 0.346 | ns | 0.2766 |
| HSR vs. OSR | -0.3674 | ns | 0.7106 |
| HSR vs. Other | -0.0493 | ns | 0.9998 |
| OSR vs. Other | 0.3181 | ns | 0.6774 |

Thapsigargin

| Comparison | Mean Diff. | Summary | Adjusted P Value |
|---|---|---|---|
| ATF6 vs. XBP1s | 0.5652 | ns | 0.2424 |
| ATF6 vs. PERK/ISR | 0.3443 | ns | 0.7605 |
| ATF6 vs. HSR | 1.7 | **** | <0.0001 |
| ATF6 vs. OSR | 1.507 | **** | <0.0001 |
| ATF6 vs. Other | 1.476 | **** | <0.0001 |
| XBP1s vs. PERK/ISR | -0.2209 | ns | 0.9522 |
| XBP1s vs. HSR | 1.135 | *** | 0.0006 |
| XBP1s vs. OSR | 0.9419 | * | 0.0284 |
| XBP1s vs. Other | 0.9107 | *** | 0.0002 |
| PERK/ISR vs. HSR | 1.356 | **** | <0.0001 |
| PERK/ISR vs. OSR | 1.163 | ** | 0.0027 |
| PERK/ISR vs. Other | 1.132 | **** | <0.0001 |
| HSR vs. OSR | -0.1931 | ns | 0.9904 |
| HSR vs. Other | -0.2243 | ns | 0.9167 |
| OSR vs. Other | -0.03122 | ns | >0.9999 |

Tunicamycin

| Comparison | Mean Diff. | Summary | Adjusted P Value |
|---|---|---|---|
| ATF6 vs. XBP1s | 0.7554 | ns | 0.0861 |
| ATF6 vs. PERK/ISR | 0.4121 | ns | 0.6888 |
| ATF6 vs. HSR | 2.009 | **** | <0.0001 |
| ATF6 vs. OSR | 1.315 | ** | 0.0020 |
| ATF6 vs. Other | 1.396 | **** | <0.0001 |
| XBP1s vs. PERK/ISR | -0.3433 | ns | 0.8096 |
| XBP1s vs. HSR | 1.253 | *** | 0.0005 |
| XBP1s vs. OSR | 0.5599 | ns | 0.5359 |
| XBP1s vs. Other | 0.641 | ns | 0.0510 |
| PERK/ISR vs. HSR | 1.597 | **** | <0.0001 |
| PERK/ISR vs. OSR | 0.9032 | ns | 0.0745 |
| PERK/ISR vs. Other | 0.9843 | *** | 0.0003 |
| HSR vs. OSR | -0.6936 | ns | 0.3409 |
| HSR vs. Other | -0.6125 | ns | 0.1277 |
| OSR vs. Other | 0.08112 | ns | 0.9998 |

CBR-470-1

| Comparison | Mean Diff. | Summary | Adjusted P Value |
|---|---|---|---|
| ATF6 vs. XBP1s | -0.06403 | ns | 0.9998 |
| ATF6 vs. PERK/ISR | 0.1011 | ns | 0.9981 |
| ATF6 vs. HSR | -0.5881 | ns | 0.1928 |
| ATF6 vs. OSR | -1.102 | ** | 0.0023 |
| ATF6 vs. Other | -0.233 | ns | 0.8402 |
| XBP1s vs. PERK/ISR | 0.1651 | ns | 0.9791 |
| XBP1s vs. HSR | -0.5241 | ns | 0.2826 |
| XBP1s vs. OSR | -1.037 | ** | 0.0040 |
| XBP1s vs. Other | -0.1689 | ns | 0.9453 |
| PERK/ISR vs. HSR | -0.6892 | ns | 0.0601 |
| PERK/ISR vs. OSR | -1.203 | *** | 0.0004 |
| PERK/ISR vs. Other | -0.3341 | ns | 0.4567 |
| HSR vs. OSR | -0.5134 | ns | 0.4963 |
| HSR vs. Other | 0.3552 | ns | 0.5156 |
| OSR vs. Other | 0.8685 | ** | 0.0065 |

Bardoxolone

| Comparison | Mean Diff. | Summary | Adjusted P Value |
|---|---|---|---|
| ATF6 vs. XBP1s | 0.0134 | ns | >0.9999 |
| ATF6 vs. PERK/ISR | -1.012 | ns | 0.2481 |
| ATF6 vs. HSR | -2.319 | **** | <0.0001 |
| ATF6 vs. OSR | -0.4154 | ns | 0.9751 |
| ATF6 vs. Other | 0.3515 | ns | 0.9368 |
| XBP1s vs. PERK/ISR | -1.026 | ns | 0.2213 |
| XBP1s vs. HSR | -2.333 | **** | <0.0001 |
| XBP1s vs. OSR | -0.4288 | ns | 0.9702 |
| XBP1s vs. Other | 0.3381 | ns | 0.9413 |
| PERK/ISR vs. HSR | -1.307 | ns | 0.0828 |
| PERK/ISR vs. OSR | 0.5971 | ns | 0.8852 |
| PERK/ISR vs. Other | 1.364 | ** | 0.0041 |
| HSR vs. OSR | 1.904 | * | 0.0143 |
| HSR vs. Other | 2.671 | **** | <0.0001 |
| OSR vs. Other | 0.7668 | ns | 0.5983 |

A3

| Comparison | Mean Diff. | Summary | Adjusted P Value |
|---|---|---|---|
| ATF6 vs. XBP1s | 0.04237 | ns | >0.9999 |
| ATF6 vs. PERK/ISR | -1.146 | ns | 0.0587 |
| ATF6 vs. HSR | -4.115 | **** | <0.0001 |
| ATF6 vs. OSR | -0.8284 | ns | 0.5256 |
| ATF6 vs. Other | -0.4475 | ns | 0.7581 |
| XBP1s vs. PERK/ISR | -1.189 | * | 0.0351 |
| XBP1s vs. HSR | -4.158 | **** | <0.0001 |
| XBP1s vs. OSR | -0.8707 | ns | 0.4458 |
| XBP1s vs. Other | -0.4899 | ns | 0.6393 |
| PERK/ISR vs. HSR | -2.969 | **** | <0.0001 |
| PERK/ISR vs. OSR | 0.318 | ns | 0.9848 |
| PERK/ISR vs. Other | 0.6989 | ns | 0.2461 |
| HSR vs. OSR | 3.287 | **** | <0.0001 |
| HSR vs. Other | 3.668 | **** | <0.0001 |
| OSR vs. Other | 0.3809 | ns | 0.9405 |

C1

| Comparison | Mean Diff. | Summary | Adjusted P Value |
|---|---|---|---|
| ATF6 vs. XBP1s | 0.112 | ns | 0.9857 |
| ATF6 vs. PERK/ISR | -0.1011 | ns | 0.9905 |
| ATF6 vs. HSR | -0.8717 | **** | <0.0001 |
| ATF6 vs. OSR | -0.2562 | ns | 0.8043 |
| ATF6 vs. Other | -0.05006 | ns | 0.9991 |
| XBP1s vs. PERK/ISR | -0.2131 | ns | 0.7880 |
| XBP1s vs. HSR | -0.9837 | **** | <0.0001 |
| XBP1s vs. OSR | -0.3681 | ns | 0.4481 |
| XBP1s vs. Other | -0.162 | ns | 0.8341 |
| PERK/ISR vs. HSR | -0.7707 | *** | 0.0003 |
| PERK/ISR vs. OSR | -0.1551 | ns | 0.9707 |
| PERK/ISR vs. Other | 0.05103 | ns | 0.9989 |
| HSR vs. OSR | 0.6156 | * | 0.0444 |
| HSR vs. Other | 0.8217 | **** | <0.0001 |
| OSR vs. Other | 0.2061 | ns | 0.8467 |

D1

| Comparison | Mean Diff. | Summary | Adjusted P Value |
|---|---|---|---|
| ATF6 vs. XBP1s | 0.2399 | ns | 0.9403 |
| ATF6 vs. PERK/ISR | -0.008735 | ns | >0.9999 |
| ATF6 vs. HSR | -1.127 | ** | 0.0010 |
| ATF6 vs. OSR | -0.1254 | ns | 0.9986 |
| ATF6 vs. Other | -0.05619 | ns | 0.9998 |
| XBP1s vs. PERK/ISR | -0.2487 | ns | 0.9187 |
| XBP1s vs. HSR | -1.367 | **** | <0.0001 |
| XBP1s vs. OSR | -0.3653 | ns | 0.8364 |
| XBP1s vs. Other | -0.2961 | ns | 0.6956 |
| PERK/ISR vs. HSR | -1.118 | *** | 0.0007 |
| PERK/ISR vs. OSR | -0.1167 | ns | 0.9989 |
| PERK/ISR vs. Other | -0.04745 | ns | 0.9999 |
| HSR vs. OSR | 1.002 | * | 0.0242 |
| HSR vs. Other | 1.071 | **** | <0.0001 |
| OSR vs. Other | 0.06923 | ns | 0.9998 |

F1

| Comparison | Mean Diff. | Summary | Adjusted P Value |
|---|---|---|---|
| ATF6 vs. XBP1s | -0.1053 | ns | 0.9987 |
| ATF6 vs. PERK/ISR | -0.6955 | ns | 0.0913 |
| ATF6 vs. HSR | -0.8594 | * | 0.0322 |
| ATF6 vs. OSR | -0.577 | ns | 0.4599 |
| ATF6 vs. Other | -0.09598 | ns | 0.9978 |
| XBP1s vs. PERK/ISR | -0.5902 | ns | 0.2073 |
| XBP1s vs. HSR | -0.7541 | ns | 0.0801 |
| XBP1s vs. OSR | -0.4717 | ns | 0.6664 |
| XBP1s vs. Other | 0.009311 | ns | >0.9999 |
| PERK/ISR vs. HSR | -0.1639 | ns | 0.9912 |
| PERK/ISR vs. OSR | 0.1185 | ns | 0.9990 |
| PERK/ISR vs. Other | 0.5995 | ns | 0.0506 |
| HSR vs. OSR | 0.2824 | ns | 0.9556 |
| HSR vs. Other | 0.7634 | * | 0.0160 |
| OSR vs. Other | 0.481 | ns | 0.5023 |

**Table S4. Whole transcriptome RNAseq of putative HSF1 activators** (see Excel Spreadsheet)

**Table S5. GO analysis of HSF1 activating compound A3** (see Excel Spreadsheet)
